# Ionic Liquid-Induced Modulation of Ubiquitin Stability: The Dominant Role of Hydrophobic Interactions

**DOI:** 10.1101/2024.12.07.627332

**Authors:** Aditya Shrivastava, Harika Kamma, Ranabir Das, Sri Rama Koti Ainavarapu

## Abstract

Despite the widespread use of imidazolium-based ionic liquids (ILs) in biotechnology, pharmaceuticals, and green chemistry, their detailed interactions with proteins, particularly affecting structural stability, remain poorly understood. This study examines the effects of ILs on ubiquitin, a thermodynamically robust protein with a β-grasp structure. We found that IL-induced destabilisation follows a consistent order with previous findings: [BF_4_]^-^ > [MeSO_4_]^-^ > [Cl]^-^ for anions and [BMIM]^+^ > [BMPyr]^+^ > [EMIM]^+^ for cations. Through pH and ionic strength-dependent studies, we observed that hydrophobic interactions predominantly influence the stability of positively charged ubiquitin, with electrostatic interactions playing a secondary role. NMR studies identified residues impacted by [BMIM][BF4]; however, mutagenesis of these residues showed minimal changes in destabilisation, suggesting a global effect. This led us to conduct a broader empirical analysis, incorporating solvent-accessible surface area evaluations, which confirmed that hydrophobic residues are the primary drivers of stability alterations in ubiquitin, with charged residues playing a minimal role. Additionally, single-molecule force spectroscopy results indicate that imidazolium ILs decrease the unfolding barrier without altering transition state structures, offering insights into protein folding dynamics. ILs appear to modulate the stability landscape of proteins by energetically and kinetically favouring the unfolded state over the folded state. These insights offer potential strategies for the selective tuning of protein stability, which could be exploited to modulate protein-protein or protein-substrate interactions in various applications of ILs.

## Introduction

Over the past two decades, ionic liquids (ILs) have emerged as versatile agents in various sectors, such as biotechnology, material science, energy solutions, green chemistry, and pharmaceutical manufacturing.^1–5^ ILs are composed of bulky and asymmetric cations and weakly coordinating anions, which due to poor packing, typically remain liquid below 100°C.^6^ Their unique structure allows for fine-tuning properties like viscosity, thermal conductivity, and hydrophobicity, thus enabling diverse applications through customized cation-anion combinations. Beyond their foundational role as ’designer solvents’ ILs have unlocked new possibilities in a myriad of applications. Their utility extends from acting as recoverable catalysts in specific chemical transformations^7^ to microemulsions for controlled synthesis of metal–organic frameworks^8^ and as antibacterial hydrogel dressing for wound healing^9^. These instances represent just a glimpse into the extensive and evolving landscape of IL applications. ILs are also recognized as green solvents due to their negligible vapor pressure^10^, a stark contrast to traditional volatile organic solvents. Nonetheless, the designation of ’green solvent’ should be approached with caution, as it does not inherently guarantee non-toxicity^11,12^. Apart from translational applications, ILs are also explored to address fundamental questions due to their interesting properties. One such area is their interaction with biological macromolecules^13–17^.

IL-protein interaction^18^ has garnered significant attention and has become a complex yet critical area of study. ILs have been used as additives for improving X-ray diffraction resolution of poorly diffracting proteins^19^, long-term protein storage^20^, protection against denaturants^21,22^, and improving the specificity^23^ and enantioselectivity^24^ of enzymes. Significant efforts have been made to understand the structure-function relationship of proteins in ILs for their rationale use. A wide range of proteins in various ILs, including azurin,^25^ BSA^26^, cytochrome c^27^, GFP^28,29^, HSA^22,30^, lipase^24,31^, lysozyme^32,33^, ribonuclease A^34^ has been studied experimentally. Parallelly, simulation-based studies^35–38^ have enhanced our understanding of these interactions at the molecular level. IL effects are often generalized via the Hofmeister series. Zhao’s efforts to quantify Hofmeister effects in ILs through viscosity B-coefficients and NMR B’ coefficients have highlighted the significant role of the Hofmeister series for anions and its ‘reverse’ for large organic cations ^39^. The organic nature and viscosity increase by long-chain cations, traditionally seen as kosmotropic, actually act as chaotropic due to less interaction with water but an enhanced interaction with hydrophobic regions of the protein^40^. Despite these insights, there are contradictions, such as with lysozyme^41^, chymotrypsin^42^, and mushroom tyrosinase^43^, where IL interactions deviate from expected patterns, underscoring the need for a deeper, case-specific understanding. Moreover, different ILs and diverse conditions used across studies further complicate comparisons and general understanding.

In this study, we have addressed this problem by systematically studying the role of imidazolium-based ILs on a protein called ubiquitin (Fig. 1a), a protein that contains an α-helix, a short piece of 3__-helix, and a mixed β-sheet containing five strands, forming a β-grasp topology. This beta-grasp fold is found in a wide range of proteins with diverse functions^44,45^. The kinetic, thermodynamic, mechanical, and various other properties of this protein have been extensively characterized^46–51^, providing a well-defined and thoroughly understood framework. Given the limited understanding of IL-protein interactions with proteins having significant beta-sheet structures^25,27,29,34,52^, ubiquitin serves as an excellent model for further investigation. By using a series of ILs (imidazolium-based) with different hydrophobicity (Fig. 1b) and counter anions, we have studied the effect of the ILs on the properties of ubiquitin. Despite ubiquitin’s positive charge, the cationic part of the ILs significantly affected its stability, suggesting the dominant role of hydrophobic interactions. Our findings advance current knowledge by showing that hydrophobic interactions do not merely influence protein stability at a local level but exert a global destabilizing effect on ubiquitin, irrespective of key residue-specific interactions. Furthermore, NMR experiments suggested local interactions of a few key residues with IL. While mutagenesis of these key residues affected protein stability, the extent of destabilization by IL remained similar with only very minute changes, suggesting the global effect on the protein rather than local residue-specific strong interactions with ubiquitin. To further elucidate this global destabilizing effect of ILs on ubiquitin, we employed single-molecule force spectroscopy (SMFS) using atomic force microscopy (AFM). SMFS is a powerful tool that allows us to study protein folding and unfolding at the single-molecule level, providing detailed insights into mechanical stability and energy landscapes. This technique has been successfully applied to various proteins, including ubiquitin, metalloproteins, and membrane proteins, enabling the characterization of unfolding forces and intermediate states. By stretching ubiquitin molecules in the presence of imidazolium-based ILs, we observed a significant reduction in the mechanical unfolding forces required. This indicates that ILs lower the kinetic barriers to unfolding, effectively making the protein’s folding energy landscape shallower. Importantly, the transition state structures remained unaltered, suggesting that ILs facilitate unfolding without causing major conformational changes at the critical unfolding points. This combination of techniques, spanning bulk to single-molecule studies, uniquely elucidates both the energetic and kinetic impacts of ILs on protein folding pathways. Our study provides a detailed analysis of the molecular effect of IL on specific physical parameters of a protein using techniques ranging from bulk to single-molecule.

**Figure 1.**
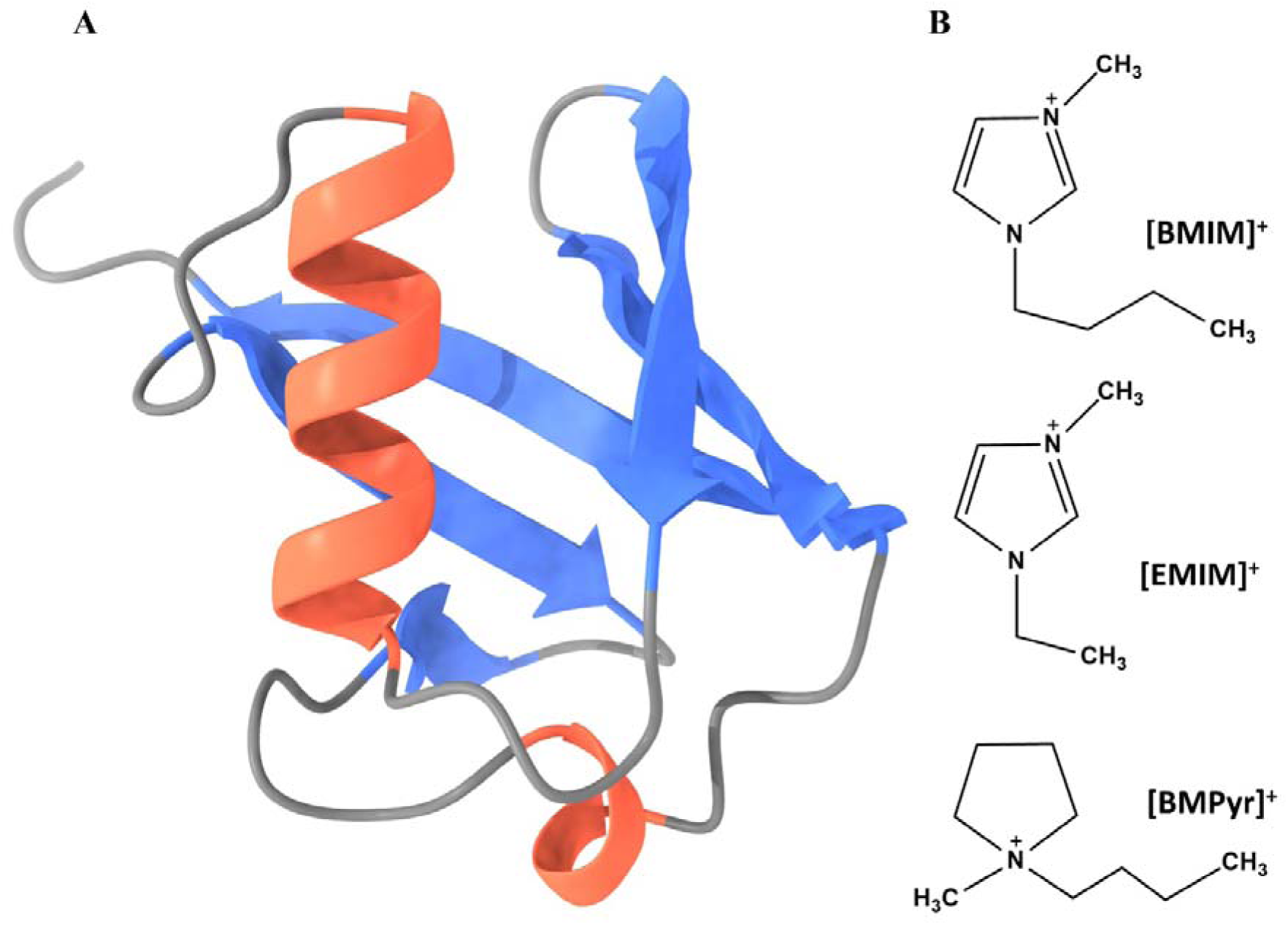
Structures of the systems used in the study. A) Cartoon diagram of ubiquitin having colour code for its secondary structure, grey – unstructured region, orange – helices, blue – β-sheets (PDB ID: 1UBQ) B) Chemical structures and formulas of Cationic part ILs used, anions are [Cl]^-^, [MeSO_4_]^-^ and, [BF_4_]^-^.

## Materials and Methods

### Gene Cloning

Alanine scanning mutagenesis of Ubiquitin F45W (UF45W) was done as described previously. PCR was performed using a high-fidelity DNA polymerase, Platinum™ SuperFi II Master Mix (Invitrogen), following the manufacturer’s instructions. (Ubq)_9_ gene was amplified by PCR from an existing plasmid (A kind gift from Prof. Julio Fernandez, pQE80L vector) and cloned into a plasmid obtained from Addgene, Spytag003-mkate2^53^ (Addgene Plasmid #133452). BamHI and KpnI restriction sites were introduced using SLIM cloning into the Spytag003-mkate2 plasmid. For, ELP_120_-SpyCatcher003 construct, BamHI restriction site was introduced into Cys-ELP(120nm)-LPETGG plasmid (Addgene Plasmid #90472), and SpyCatcher003 S49C (Addgene Plasmid #133448) was cloned into it using BamHI and XhoI restriction sites, followed by SpyCatcher003 C49S mutation. Sequencing of plasmids was done by Sanger Sequencing (Eurofins India, Bengaluru). Sequences of used protein constructs are provided in the supplementary information.

### Protein Expression and Purification

The plasmids containing ubiquitin (UF45W, Spytag003-(Ubq)_9_) were transformed into Rosetta 2(DE3) ΔElaD *E. coli* competent cells (Gifted by Prof. Ronald T. Hay, University of Dundee), whereas ELP_120_-SpyCatcher003 S49C^53^ was transformed into BL21(DE3) *E. coli* competent cells. A single colony was picked and inoculated into LB Media at 37 °C, 200 r.p.m overnight. The overnight culture was diluted into 1 L of LB media and kept at 37, 200 r.p.m. The protein expression was induced by adding IPTG (final concentration of 1mM) after the OD_600nm_ of the secondary culture reached 0.8-1.0. The cells from the secondary culture were then pelleted down by centrifugation at 6000 r.p.m. for 15 minutes at 4°C. The pellet was resuspended in lysis buffer (50 mM Phosphate buffer, pH 8.0, 300 mM NaCl, 10 mg Lysozyme, 200 µL of Triton-X100, and cOmplete Protease Inhibitor Cocktail tablet) and lysed by sonication. The lysate was then centrifuged at 18000 r.p.m. for 20 minutes at 4°C to separate the soluble and insoluble fractions. The soluble fraction was used for further purification.

For protein purification, a Ni-NTA (Nickel Nitrilotriacetic acid) affinity chromatography was performed. The soluble fraction was loaded onto a Ni-NTA column pre-equilibrated with PBS buffer (50 mM Phosphate buffer, 150 mM NaCl). The column was then washed with the same buffer to remove any unbound proteins. The proteins of interest were then eluted using PBS buffer (50 mM Phosphate buffer, 150 mM NaCl, 300 mM Imidazole).

The eluted fractions were then concentrated using Amicon Ultra-15 Centrifugal Filter Units. The concentrated proteins were further purified by size-exclusion chromatography using a HiLoad^TM^ 16/600 Superdex^TM^ 200 pg for Spy3-Ubq_9_ and a HiLoad^TM^ 16/600 Superdex^TM^ 75 pg column for smaller proteins. The proteins were eluted in PBS buffer (50 mM Phosphate buffer, 150 mM NaCl, 200 mg/L NaN_3_, pH 7.4). The purity of the proteins was assessed by SDS-PAGE. Prior to further studies, UF45W was dialyzed against 50 mM citrate buffer and 150 mM NaCl at pH 3.0/3.5/4.0 to prepare for further experiments.

This comprehensive purification protocol allowed us to obtain highly pure proteins for further studies.

### Chemicals used

3-Aminopropyl(diethoxy)methylsilane (97%), [BMIM][BF_4_] (≥97.0% (HPLC)), [BMIM][Cl] (≥99.0% (HPLC)), [BMIM][MeSO_4_] (≥97.0% (HPLC)), [EMIM][BF_4_] (≥97.0% (HPLC)), [BMPyr][Cl] (≥99%(T)), [EMIM][Cl] (>99%), [EMIM][MeSO_4_] (≥98.0% (HPLC)), Sodium Citrate, Sodium Borate, Citric Acid were procured from Sigma Aldrich Ltd. Maleimide-PEG_2_-NHS Ester was procured from TCI (India).

### Near-UV Circular Dichroism Experiments

Temperature-dependent near-UV circular dichroism (CD) studies were carried out in different ionic liquids by a JASCO J-1500 (JASCO Corp., Japan) CD spectrophotometer. CD spectra for the tertiary structure region (>250 nm) were recorded using ∼125 μM UF45W in a quartz cuvette (Starna Scientific, 53/Q/SOG/1) of 1 cm path length. For analysis, an average of ellipticity θ from 265-269 nm was taken and plotted against the temperature and fitted using a two-state model^54^ as described below –

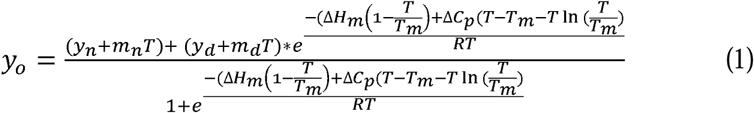

where y_o_ = ellipticity, y_n_ and y_d_ represent the intercepts, and m_n_ and m_d_ are the slopes of native and unfolded baselines. T_m_, ΔH_m_ and ΔC_p_ are melting temperature, enthalpy change, and heat capacity change on unfolding at T_m_ .

### Backbone assignment

Ubiquitin was prepared in 50mM sodium citrate (pH 3.5) with 150mM NaCl. Standard triple resonance experiments, HNCACB and CBCA(CO)NH, were performed to assign backbone amides in Ub 15N-1H HSQC spectra obtained in the above buffer conditions in NMRFAM Sparky.

### NMR Titration

Both protein and ligand stocks were prepared in 50mM sodium citrate (pH 3.5) with 150mM NaCl for Ub: BMIM-BF4 titration. Ubiquitin was titrated with 2M BMIM-BF4 up to 640mM (=1:3616 ratio).^15^N-^1^H HSQC spectrum was recorded for each titration point at 298K on 600 MHz Bruker NMR Spectrometer, followed by processing in NMRPipe and peak assignment in NMRFAM Sparky. Chemical shift perturbation (CSP) was calculated for each residue after every titration point using the equation, 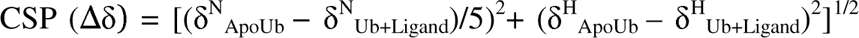, where δ^H^ and δ^N^ denote amide hydrogen and nitrogen chemical shifts, respectively, in the presence and absence of ligand. For determining K_d_, calculated CSPs were fit into a 1:1 protein: ligand model using the equation, Δδ_obs_ = Δδ_max_{([P]_t_+[L]_t_+K_d_) - √([P]_t_+[L]_t_+K_d_)^2^-4[P]_t_[L]_t_}/2[P]_t_, where Δδ_max_ represents CSP calculated for the last titration point, and [P]_t_ and [L]_t_ are protein and ligand concentrations at a titration point.

### SASA Ranked CSP Analysis (SRCA)

The SRCA method applies a sliding window approach to analyze residues with similar solvent-accessible surface area (SASA), minimizing biases from residue type frequency and enabling systematic comparisons of CSP patterns across similarly exposed regions. To calculate residue-wise solvent-accessible surface areas (SASA) of ubiquitin we used FreeSASA^55^. Residues were categorized into four types—Hydrophobic, Polar charged, Polar noncharged, and Glycine—based on their physicochemical properties. The analysis was performed on ubiquitin (PDB: 1UBQ). A CSV file was generated containing residue identifiers (e.g., M1, Q2), SASA values, and residue categories. CSP data from the final titration point (640mM) was then incorporated into this file, resulting in a final CSV containing the columns Residue, SASA, CSPs, and Type of residue. The Python script used for this calculation is provided in Supplementary Script 1. Using the final CSV file, a sliding window approach was applied to analyze residue chemical shift perturbations (CSPs). Analyses were performed using window sizes of 3, 4, and 5 residues. For each window, subsets containing at least three different residue types were identified. For each subset, the residue with the highest CSP value was recorded, along with its residue category. The results of the sliding window analyses are summarized in Table S1, and the Python script used for this analysis is provided in Supplementary Script 2.

### AFM Experiments

Glass coverslips were cleaned and activated using hot chromic acid (80 °C) for 2 hours and rinsed with acetone and water. We dried them under nitrogen stream, and they were further activated using oxygen plasma for 30 minutes. We incubated them with 1% APDMS in ethanol for 1 hour followed by washing with ethanol and gentle drying with N_2_. We baked the coverslips for 1 hour at 80°C followed by incubation with a solution of 1 mg/mL maleimide-PEG_2_-NHS ester in a mixture of 50% DMSO and 50% 50 mM sodium borate buffer (pH 8) for 3 hours at RT. We rinsed them using DMSO and isopropanol, respectively and, dried them under an N_2_ stream, stored them at -20 °C till their further use. Covalent functionalisation was confirmed using confocal microscopy (Fig. S1). Before the AFM experiment, coverslips were incubated with 30 μM ELP_120_-SpyCatcher003 for 3 hours at room temperature. We rinsed them using a PBS buffer. Coverslip tethered with ELP_120_-Spycatcher003 were incubated with 10 μM SpyTag003-(Ubq)_9_ for 1 hour and rinsed with PBS. We finally incubated the coverslip with IL solution for 30 mins. Constant velocity experiments were performed at 100, 400, 1000, 2000, and 4000 nm/s speeds using a home-built AFM setup. A schematic of the AFM experimental setup is shown in Supplementary Fig. SX.

### AFM Data Analysis

For the analysis of AFM data, only force-versus-extension (FX) profiles displaying a minimum of three unfolding force peaks were selected. These profiles were analysed using the worm-like chain (WLC) model to describe polymer elasticity (Eq. 2). The model equation is expressed as follows, where F represents the applied force, x is the molecular extension, p the persistence length, L_c_ the contour length, k_B_ Boltzmann’s constant, and T the temperature set at 298 K. To achieve the best fit for all unfolding peaks within an FX trace, the persistence length parameter was adjusted between 0.3 and 0.6 nm.

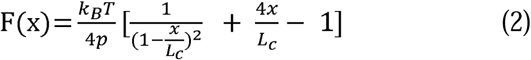

The calculation of the distance to the transition state (Δx_u_) and the spontaneous unfolding rate constant (k^O^) employed using Monte Carlo simulations. The activation energy (ΔG^‡^) was derived using the Arrhenius equation (Eq. 3). Protein’s spring constant (*k*_s_) in the direction of N- to C-terminus pulling was determined via harmonic approximation (Eq. 4) ^56^. Where, k_A_ is the Arrhenius frequency factor. For protein dynamics, k_A_ has a value of 10^9^ s^−1^ ^57^.

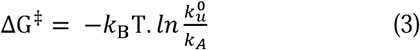

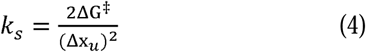

## Results

### Thermal Stability of Ubiquitin in Ionic Liquids

To assess the thermal stability of ubiquitin (UF45W, Tryptophan variant of ubiquitin^58^) in different ionic liquid environments, we performed temperature-dependent near-UV circular dichroism (CD) experiments of the protein at pH 3.5 (Fig. 2A). The melting temperatures (T_m_) were determined by monitoring the change in ellipticity at 267 nm as a function of temperature (Fig 2B). The T_m_ (351.7 K) obtained for the protein from CD was found to be in good agreement with the T_m_ (352.7 K) obtained from differential scanning calorimetry (DSC) (Fig. S2), suggesting that the protein undergoes a cooperative unfolding event and no observable intermediate exists, as expected^59^. Our findings also indicated that the tertiary structure of UF45W was retained across all conditions tested. Slight decrements were observed in the near-UV CD spectrum while ubiquitin was subjected to various ILs. However, the impact of ionic liquids on UF45W thermal stability was significant. We quantified this by monitoring the change in T_m_ at two different concentrations of ILs (0.5 M and 1 M, respectively). The result for 1M is shown in Fig. 2C (0.5 M in Fig. S3). Further, to elucidate the effects of the ionic liquid constituents on protein stability, we calculated the ΔT_m_, which is defined as

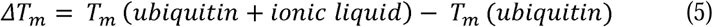

**Figure 2.**
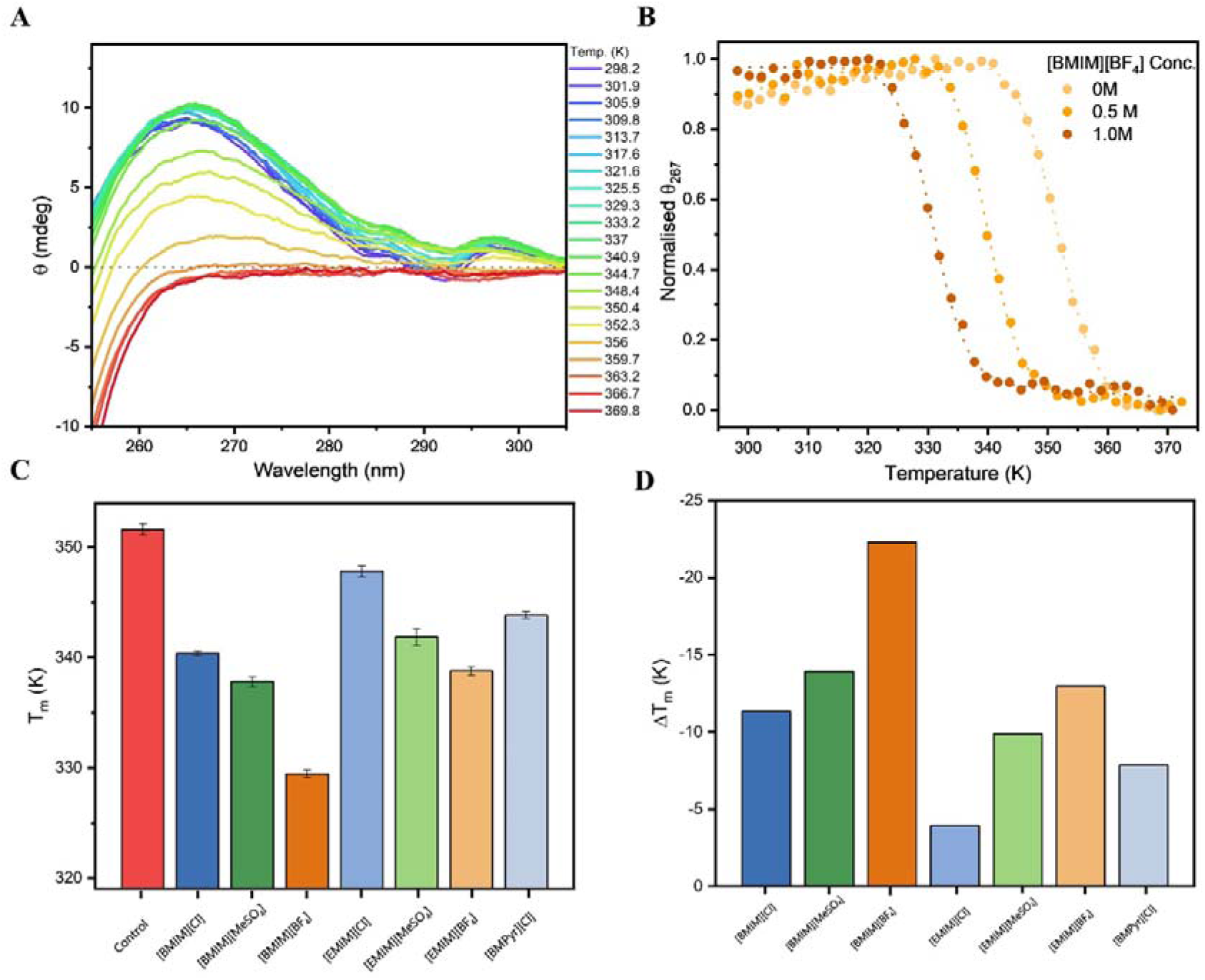
Analysis of Ubiquitin F45W Thermal Stability in various ILs. A) Temperature-dependent Near-UV CD spectra of ubiquitin F45W. B) Ellipticity at 267 nm plotted against temperature reveals T_m_ variations, signifying alterations in thermal stability due to ionic liquid interactions. C) Bar graph representing T_m_ analysis in different ionic liquids at 1 M concentration, demonstrating the influence of ionic environments on protein stability D) Bar graph representing ΔT_m_ quantification illustrates the varying degrees of ubiquitin destabilisation across different ionic liquid compositions at 1M, highlighting the differential impacts of cation and anion constituents on protein stability.

-We arranged the cations and anions based on their order of destabilising impact (Fig. 2D). The extent of destabilisation as probed by a change in the melting temperature (|ΔTm|) by cations after keeping anion (Cl^-^) constant: [BMIM]^+^ (11.1 K) > [BMPyr]^+^ (7.8 K) > [EMIM]^+^ (3.7K), and by anions after keeping cation (BMIM^+^) constant: [BF_4_]^-^ (22.0K) > [MeSO_4_]^-^ (13.9) > [Cl]^-^ (11.9K).

### pH and Ionic Strength-Dependent Stability of Ubiquitin in Ionic Liquids

To further understand the effect of ionic strength and pH on ionic liquid-mediated ubiquitin destabilisation, we performed additional temperature denaturation measurements in CD. Especially imidazolium-based ionic liquids ([BMIM][BF4], [BMIM]Cl, and [EMIM][BF4]) were chosen at varying pH levels (3.0, 3.5, and 4.0) and ionic strengths (150 mM NaCl and 1M NaCl). As the pH increased from 3.0 to 4.0, the net positive of the UF45W charge decreased by ∼6 units due to the deprotonation of acidic side chains (Fig. S4)^60–62^. The positive charge on the imidazolium ring, however, remained constant across these various ILs. The T_m_ of ubiquitin was measured in [BMIM][BF4] across a pH range of 3.0 to 4.0 (Fig. 3A). The absolute T_m_ values of ubiquitin increased with an increase in pH. We measured the ΔTm value as a function of pH. The ΔTm value also increased as a function of pH (Fig. 3B). We extended this to other ionic liquids and determined the corresponding ΔT_m_’s. At pH 4.0, we observed a similar increase in destabilisation across all three ILs (ΔTm), irrespective of their alkyl side chain length (BMIM and EMIM) or nature of counter anion (BF_4_^-^ and Cl^-^). Although the absolute decrease in T_m_ with different IL followed the same order irrespective of pH, the ΔTm values suggested the involvement of another destabilisation factor apart from hydrophobicity (most likely a coulombic interaction).

**Figure 3.**
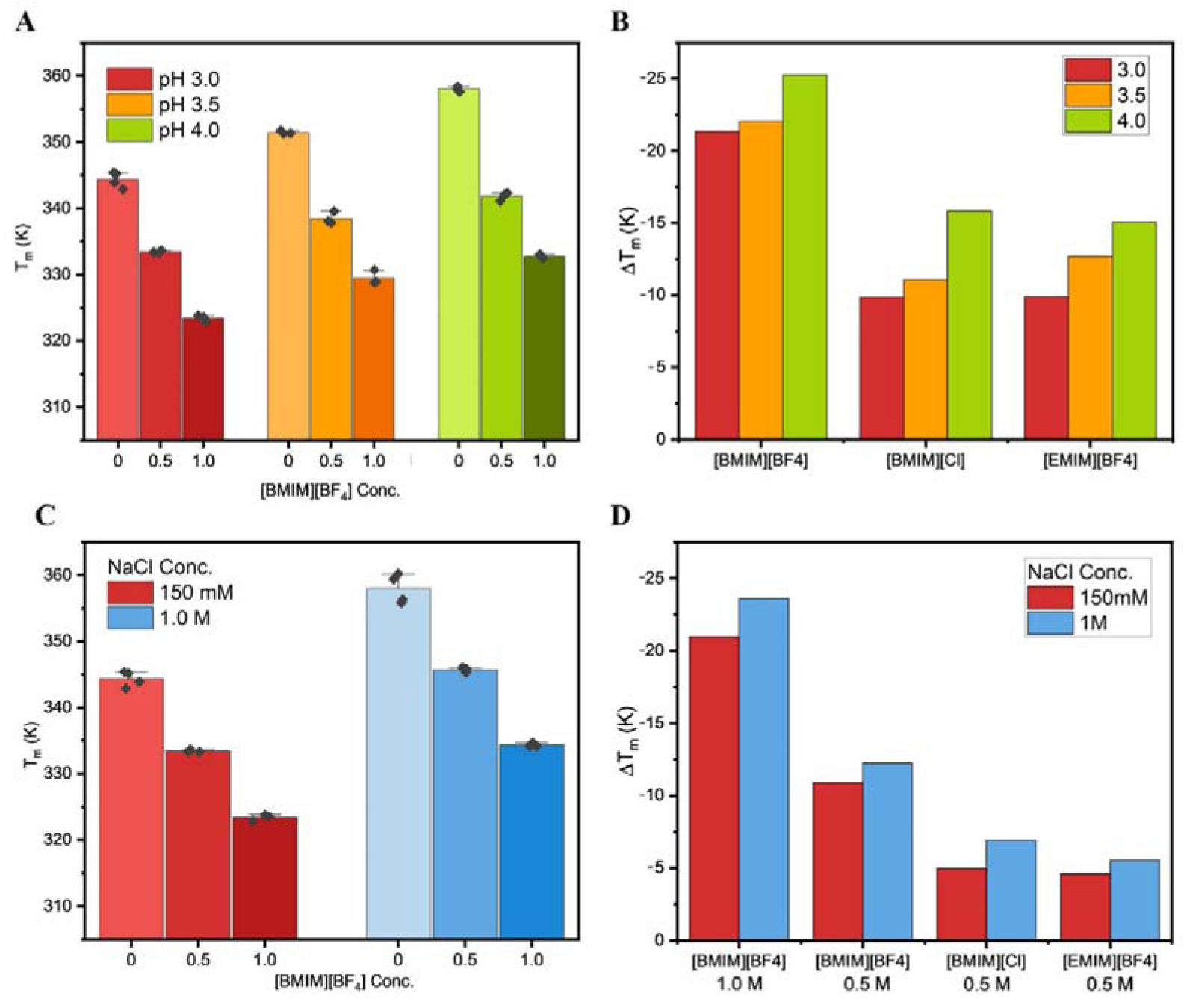
Influence of pH and ionic strength on ubiquitin stability in imidazolium-based ILs A) Bar graph representing the effect of pH on T_m_ of ubiquitin. B) Bar graph representing the change in ΔT_m_ across different pH levels for ubiquitin in 1M [BMIM][BF4], [BMIM]Cl, and [EMIM][BF4], illustrating the incremental destabilisation of the protein structure as pH increases. This effect is noted regardless of the ionic liquid’s counter anion and length of the alkyl side chain of the cation, suggesting a common destabilising interaction that intensifies as the protein’s net positive charge decreases. C) Bar graph representing the effect of ionic strength on ubiquitin’s T_m_ at pH 3.0, exploring how varying concentrations of NaCl influence protein stability at various concentrations D) Bar graph representing the minor differences in ΔT_m_ due to varying ionic strengths, illustrating the nature of interactions between ubiquitin and imidazolium-based ionic liquids. A lesser extent of destabilisation at high ionic strength suggests a minor role of coulombic interactions at pH 3.0.

To test that out, we also measured the effect of ionic strength on ubiquitin stability at pH 3.0. High ionic strength (1M NaCl) screens the repulsion among positively charged groups on the surface of ubiquitin, especially at pH 3, leading to stabilisation (Fig. 3C). The change in magnitude of destabilisation induced by Ionic Liquids at high ionic strength was equivalent to the pH-dependent destabilisation (LJT_m_ 20.9 at pH 3, 25.2 at pH 4, and 23.6 at pH 3, 1 M NaCl). This indicated a dual role of both Coulombic interaction and hydrophobicity at pH 3.0 (Fig 3D).

### NMR Titration Analysis Identifies Key Residues in Ubiquitin Interacting with [BMIM][BF4]

To identify individual residues in ubiquitin and their nature of interaction with imidazolium-based ILs in a more quantifiable manner, we performed an NMR titration analysis of ubiquitin using [BMIM][BF4]. This choice was guided by its pronounced impact on the thermal stability of ubiquitin; [BMIM][BF4] destabilised ubiquitin by 22K (at pH 3.5), the most significant decrease observed among all the ILs we screened. We observed significant chemical shift perturbations (CSPs) that provided insights into the residues involved in the interaction with the ionic liquid. The 15N-1H HSQC spectra at different molar concentrations (Fig. 4A) revealed distinct changes in the resonances of certain residues. We observed the most significant perturbations (Mean + 2SD) in residues G47, E16, and L71 (Fig. 4B), indicating their direct involvement in interactions with [BMIM][BF4]. Additionally, residues Q2, L8, I13, K33, and L73 also showed notable deviations (Mean + SD), suggesting a secondary tier of influence from the IL.

**Figure 4.**
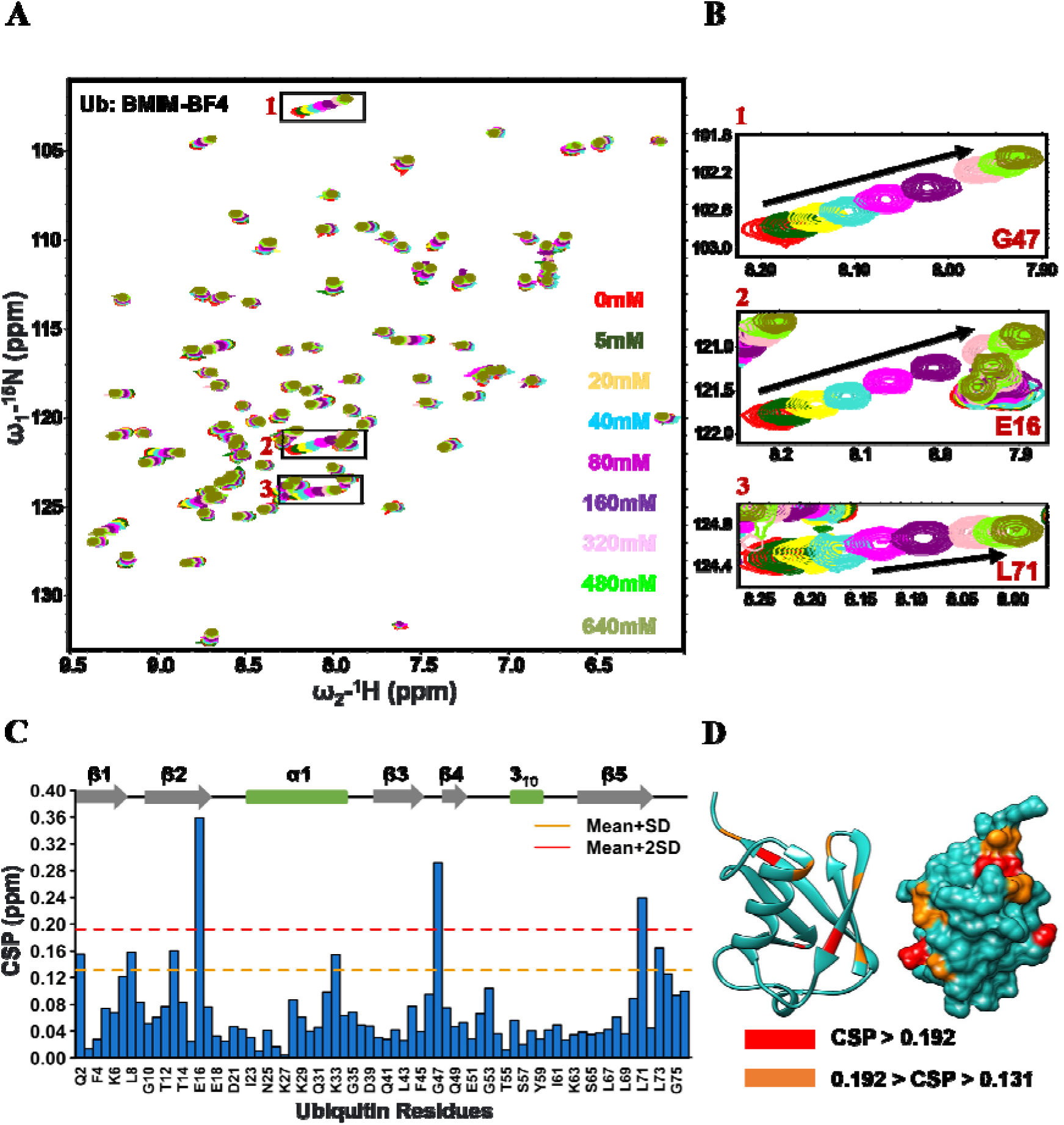
NMR Titration analysis for Ubiquitin in BMIM-BF4. A) Overlay of ^15^N-^1^H HSQC spectra at different molar concentrations. B) Zoomed-in view of the resonances of G47, E16, and L71 showing significant perturbation. C) Chemical shift perturbations (CSPs) quantified and plotted against Ubiquitin residues. Dashed lines represent the thresholds beyond which CSPs were considered significant. The orange line marks residues with CSPs above Mean + SD. The red line marks residues with CSPs above Mean + 2SD. D) Residues showing significant perturbations beyond these thresholds were mapped onto ubiquitin using the same colour code (as described in C).

We quantified the CSPs and plotted them against the ubiquitin residues (Fig. 4C). We set thresholds to identify significant CSPs: residues with CSPs above the mean plus one standard deviation (SD) (marked by an orange line) and those with CSPs above the mean plus two SDs (marked by a red line). Residues showing significant perturbations beyond these thresholds were mapped onto the structure of ubiquitin (Fig. 4D) using the same colour code as in figure 4C. This visual representation highlighted the spatial distribution of these key residues on the ubiquitin molecule, providing a clearer understanding of how ubiquitin interacts with [BMIM][BF4]. The perturbation is significant at the surface of β1, β2, β4, and the C-terminal end of β5. The alpha helix is not perturbed by [BMIM][BF4].

### Mutagenesis Studies Reveal Subtle Changes in Ubiquitin Stability in the Presence of [BMIM][BF4]

We probed the role of the three residues that showed the highest CSP using site-directed mutagenesis. We created three single mutants (G47A, E16A, and L71A). The mutation of these residues allowed us to assess their individual contributions to the interaction with [BMIM][BF4]. We also included a double salt bridge null mutant (K11A/E34L/K27A/D52L) termed as DS which we previously had shown to be of similar chemical stability as of wild-type ubiquitin.

For L71A, we reduced the size of the hydrophobic side chain by substituting the larger leucine residue with the smaller alanine. In the case of G47A, we introduced a slight increase in hydrophobicity by replacing the unique glycine, which lacks a side chain, with alanine. With E16A, we removed the polar, charged part of the glutamate residue and reduced the side chain’s hydrophobic length by substituting it with alanine.

The thermal stability of these mutants, represented by their melting temperatures (Tm), was compared both in the absence and presence of [BMIM][BF4]. In the absence of the ionic liquid, all single mutants displayed a decreased T_m_ relative to the wild type (WT), indicating an inherent destabilisation due to these mutations, while the double salt-bridge null mutants showed an increment in T_m_. The order of stability, from most to least, was DS ≥ WT > E16A ≈ L71A > G47A (Fig 5A). Upon adding [BMIM][BF4], all mutants showed destabilisation. To quantify the minor changes in stability due to the ionic liquid for each mutant relative to the WT, we calculated the ΔΔT_m_ (Fig. 5B). It is defined as

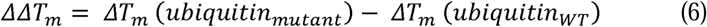

**Figure 5.**
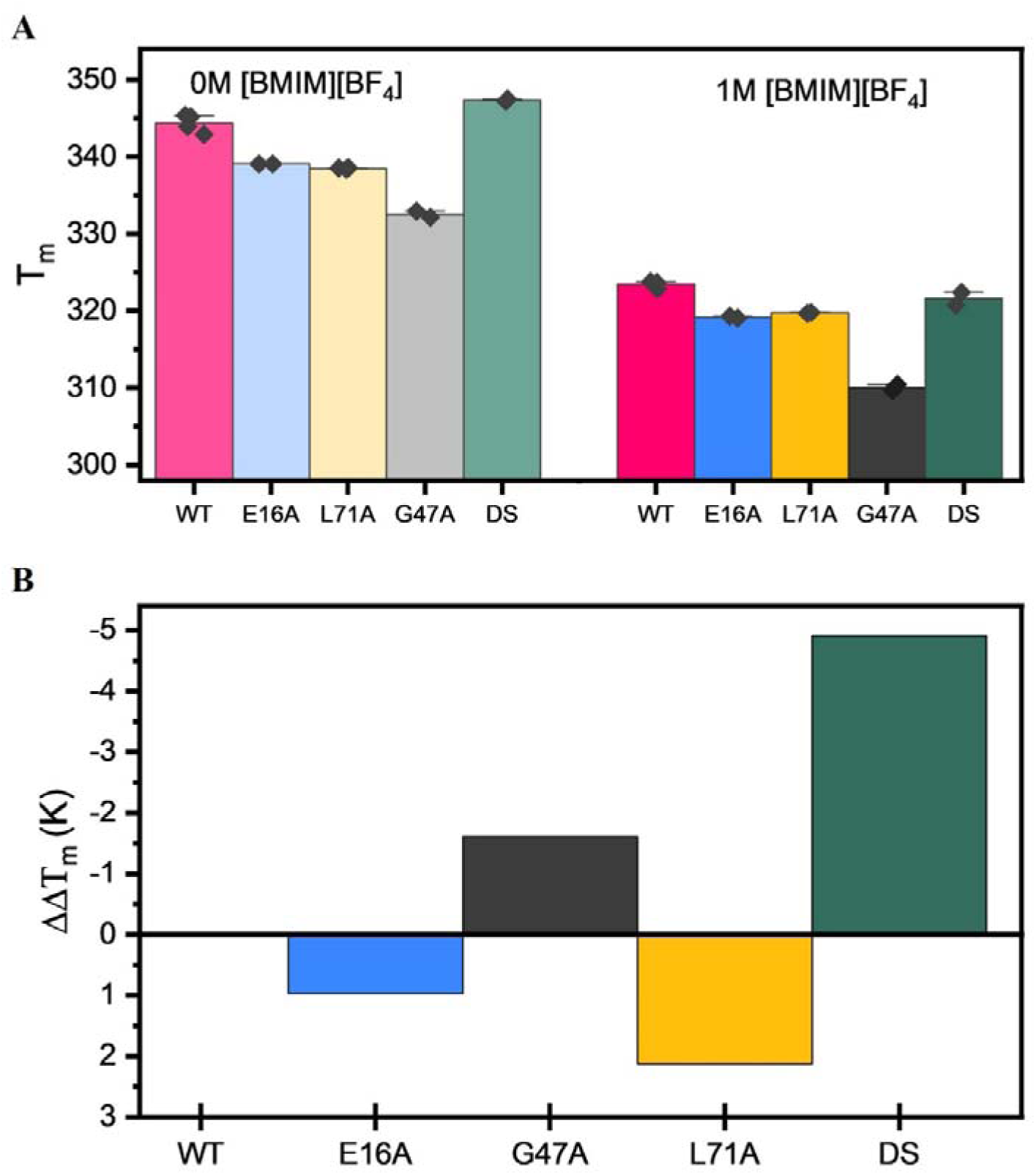
Impact of mutagenesis of some of the key residues on the ubiquitin’s stability in 1M [BMIM][BF4] A) Bar graph representing the effect of various mutagenesis on T_m_ of ubiquitin B) Bar graph depicting the changes in ΔΔT_m_ across various mutants indicating slight variations in stability. (here, DS = K11A/E34L/K27A/D52L)

If the mutation reduces IL binding and destabilisation, the ΔΔT_m_ value should be positive. The E16A (ΔΔT_m_ = 1.0) and L71A (ΔΔT_m_ = 2.1) resulted in slightly less destabilisation [BMIM][BF4] compared to WT, whereas the G47A (ΔΔT_m_ = -1.6) mutation led to a marginally increased destabilisation. DS (ΔΔT_m_ = -4.9) showed a significantly higher increase in destabilisation.

### SASA Ranked CSP Analysis (SRCA) for Global Analysis of Ubiquitin-IL Interactions

In our study, we introduced a unique empirical approach, the SASA Ranked CSP Analysis (SRCA), to probe the global interactions between ubiquitin and the ionic liquid [BMIM][BF4]. Traditional CSP analysis identifies perturbed residues but lacks structural context, often solvent accessibility and residue type distribution. The SRCA method addresses these gaps by integrating CSP data with SASA rankings to correlate solvent-accessible surface area (SASA) and chemical shift perturbations (CSPs) of protein residues. The residues were categorized into four types—hydrophobic, glycine, polar noncharged, and polar charged—and arranged in order of decreasing SASA. This arrangement was based on the assumption that exposed residues have a higher probability of interacting with the IL. In this arrangement, for each moving set of five consecutive residues with similar SASAs, we identified the residue type with the highest CSP. This analysis yielded 41 sets of five consecutive residues, each containing at least three different residue types (additional analysis considering moving sets of 4 and 6 are provided in Table S1). This approach allowed us to have a meaningful comparison, the results revealed a distinct pattern: hydrophobic residues had the highest CSP in 15 out of 41 sets, followed by glycine residues in 12 sets, polar non-charged residues in 11 sets, and polar charged residues in 3 sets. This suggested that the environment of the hydrophobic residues was more prone to perturbation by the IL [BMIM][BF4] than that of the polar charged residues.

However, it is important to note that these results do not simply reflect the overall distribution of residue types in ubiquitin. In the 41 sets, hydrophobic residues appeared 39 times, polar charged residues 32 times, polar non-charged residues 38 times, and glycine residues 20 times. Thus, the overrepresentation of hydrophobic and glycine residues and the underrepresentation of polar charged residues in the sets with the highest CSPs cannot be attributed to their overall frequency in the protein.

### Mechanical Stability of Ubiquitin in the Presence of [BMIM][BF4] at Physiological pH

To investigate ubiquitin’s behaviour at physiological pH of 7.4, where its T_m_ exceeds 100 °C^59^, we used atomic force microscopy (AFM)-based single-molecule force spectroscopy (SMFS). This approach allowed us to study the unfolding behaviour of ubiquitin in the presence of [BMIM][BF4] at pH 7.4, bypassing the limitations of traditional thermal stability assessments. Moreover, SMFS enabled us to explore changes in the folding energy landscape of ubiquitin in the presence of [BMIM][BF4]. The use of force spectroscopy is particularly relevant in biotechnological contexts, such as in bioreactors or during vigorous mixing, where proteins often encounter shear forces^63^. We obtained force-extension (FX) curves having sawtooth-like pattern (Fig. 7A). The unfolding forces observed for ubiquitin in [BMIM][BF4] exhibited a noticeable decrease at all pulling velocities (Fig. 7B). This reduction underscores a fundamental shift in the mechanical resistance of ubiquitin, suggesting that [BMIM][BF4] also altered the kinetic barriers encountered along the unfolding pathway, keeping the contour length (ΔL_c_) of ubiquitin approximately similar (Fig. S5) with that measured in aqueous environments. This constancy indicates that while [BMIM][BF4] affects the forces required to initiate unfolding, it does not alter the ultimate extension of the protein, preserving its elongation characteristics. To further dissect the mechanistic underpinnings of these observations, we performed Monte Carlo simulations. Remarkably, the unfolding rate constant (k_u_) was found to increase by a factor of ∼2 in the presence of 1M [BMIM][BF4], while the distance to the transition state (x_u_) showed no change (Table 1). This suggested that [BMIM][BF4] primarily influences the energetic landscape by lowering the unfolding energy barrier, thereby accelerating the unfolding process without significantly affecting the structural configuration at the transition state.

**Table 1.** Comparison of mechanical properties of ubiquitin in absence and presence of 1M [BMIM][BF4].

| Quantities | native condition (Control) | 1M [BMIM][BF<sub>4</sub>] |
| --- | --- | --- |
| unfolding force (pN) | 204±30 | 186±31 |
| $\Delta L_c$ (nm) | 23.1±0.8 | 23.5±0.6 |
| $\Delta x_u$ (nm) | 0.2 | 0.2 |
| $k_u^\circ$ (s<sup>-1</sup>) | 0.0070 | 0.0137 |
| $\Delta G^\ddagger$ ($k_B T$) | 25.7 | 25.0 |
| $k_s$ (N/m) | 5.3 | 5.1 |

## Discussion

### Ubiquitin Stability Across Varying Ionic Liquid Conditions

At first, by taking a series of different ILs, we ventured onto understanding the effects of charge and hydrophobicity on the thermal stability of ubiquitin. A lack of intermediates, and stable tertiary structure indicated a two-state unfolding process in presence of IL. We deliberately altered the pH of the solution to a lower value (pH 3.5) to obtain a reasonable Tm value^59^. All the ILs decreased the T_m_ in an order of [BMIM]^+^ > [BMPyr]^+^ > [EMIM]^+^ for the cations, and [BF_4_]^-^ > [MeSO_4_]^-^ > [Cl]^-^ for anions.

The cationic order of aligns with observations in RNAse I study^34^, where cations follow the reverse Hofmeister series correlated (i.e., more kosmotropic cations destabilize proteins) and correlates with hydrophobicity. As alkyl chain length increases, these cations often penetrate protein surfaces more effectively, leading to greater destabilization. This behavior has been observed in several proteins, including azurin ^25^ and human serum albumin (HSA)^64^, where a combination of CW EPR and DEER spectroscopy revealed that [BMIM]^+^ caused significant structural perturbation and unfolding compared to shorter alkyl chain analogs like [EMIM]^+^^25^. Additionally, the difference between [BMIM]^+^ and [BMPyr]^+^ is due to the positive charge being delocalized over the ring in [BMIM]^+^, making it more hydrophobic than [BMPyr]^+^. This trend has also been observed in GFP^29^, highlighting the universal impact of hydrophobic interactions facilitated by longer alkyl chains and aromaticity. However, an intriguing deviation is noted in the anionic destabilisation order [BF_4_]^-^ > [MeSO4]^-^ > [Cl]^-^, where [MeSO4]^-^ is typically more kosmotropic than Cl^-^ according to the Hofmeister series^65,66^.

However, on increasing the pH slightly (pH 4), the extent of destabilisation increased with a reduced difference among different ILs. Higher pH results in decrease in net positive charge in ubiquitin resulting in protein stability (increased T_m_, Fig. 3A). This can also lead to a decreased repulsive interaction with the positively charged ILs resulting in higher approachability and more destabilisation. We hypothesize the decrease in ΔT_m_ across different IL stems from a steric hindrance where the lesser bulky (also less hydrophobic) IL can approach ubiquitin more. Thus, this suggested a direct dependence of hydrophobicity on the thermal destabilisation of ubiquitin. To test out this hypothesis further, we performed our experiments at a higher salt concentration (1M NaCl) over physiological 150 mM salt. We found a similar extent of higher destabilisation as that of pH 4. Additionally, our observations indicate imidazolium-based ILs reduce ubiquitin aggregation during the thermal denaturation (Fig. S6), as explained, potentially disrupting hydrophobic interactions between aggregates, as reported earlier for RNAse I^67^. This signified that hydrophobicity was a major contributing factor in determining the extent of destabilisation of ubiquitin by the different ILs.

### Subtle Residue Specific Effect of [BMIM][BF_4_] on Ubiquitin

Having established that by varying the hydrophobicity and aromaticity of cations and anions, and by changing the solution conditions such as changing pH and ionic strength, we further delved deeper to understand the effect of IL from a molecular perspective. CSP of individual residues of ubiquitin was measured using HSQC NMR. To understand residue-specific effects of IL on ubiquitin, we chose [BMIM][BF4] which showed the maximum destabilisation. ΔCSP values of ubiquitin calculated from different experimental conditions (with and without [BMIM][BF4]) pointed three residues to be maximally affected (with > 2 SD). Those were E16, G47, and L71 in an order E16 > G47 > L71. Moreover, other four residues in the order of L72 > I13 > L8 > Q8 also showed significant perturbation (1SD < CSP <2SD). In a previous report of GB1, a β-grasp protein (same as ubiquitin) ^68^ where specific nonpolar residues such as V21 and V29, were impacted by [BMIM][Br] interactions. In contrast, GFP, a beta-barrel protein^29^, where no specific preference for residue type or location was observed in response to different [BMIM] based ILs, showing a diffuse pattern of interaction with ILs. On the other hand, Lipase A, an α/β hydrolase^52^, exhibited a distinctly localized interaction, with significant perturbations concentrated in a patch of residues including the catalytic H156 and G158. This suggests a more targeted interaction pattern, presumably driven by hydrophobic interactions with the butyl chain of [BMIM]. These observations highlight the complexity and heterogeneity of protein IL-protein interactions. Although from the previous bulk measurements, we found the destabilising factors not to be electrostatic, in order to understand the role of each of these three amino acids (E16, G47, L71) in the ubiquitin’s stability, we performed an alanine scan site-directed mutagenesis. Site directed replacement of all these residues with alanine resulted in a decreased stability (governed by a concomitant decrease in T_m_). However, on calculating the ΔΔTm as stated in equation 2, interesting trends were observed. For both E16A and L71A, the values were positive. This indicated that the destabilisation in the mutated ubiquitin was lower than the WT. We hypothesised that the decrease in hydrophobicity moving from leucine to alanine, and glutamic acid to alanine could be the reason behind lower destabilisation. Although E is charged, the bulky side chain compared to alanine could be the reason. Therefore, we expected that by increasing the hydrophobicity of the side chains, the IL-induced destabilisation in ubiquitin should be more (or ΔΔTm should be negative). This is exactly what we observed with the G47A mutant (Fig. 5B). However, changes in destabilisation by [BMIM][BF_4_] due to the replacement of key residues were subtle (|ΔΔTm|<2K), indicating a non-specific or global phenomenon rather than residue-specific interactions. Similarly, NMR studies on the small alpha-helical protein Im7 have shown that both specific ion-protein interactions and non-specific effects contribute to destabilization by imidazolium-based ILs^69^. When both cations and anions interact strongly with the protein, the net result resembles a non-specific interaction, ultimately leading to unfolding. After our mutagenesis analysis based on HSQC NMR did not reveal significant destabilisation. However, a cluster of mutations involving the residues with the highest CSP values might nullify the IL effect. To this end, we made three mutations together (E16A/ /G47A/L71A); however, the resulting protein is drastically less stable, so it couldn’t not be explored further. Also, these residues might be very important for the folding and stability of the protein. Moreover, the fit of NMR titration data using a 1:1 protein-ligand model yielded an average K_d_ of 311±8 mM (Fig. S7). This notably high K_d_ suggests the individual IL-residue interactions between ubiquitin and [BMIM][BF_4_] are weak, but distributed over the entire protein surface, corroborating our findings of minimal changes in ΔΔT_m_ and supporting a non-specific interaction mechanism.

To support these findings from ionic strength experiments that the ubiquitin-IL hydrophobic interactions play a dominant role, we have made another mutant of ubiquitin, K11A/E34L/K27A/D52L, this mutant has increased the hydrophobic groups. Interestingly, this mutant is as stable as the WT protein and allowed us to test this hypothesis.^51^ Despite low CSP values for these residues, this mutant showed significantly higher destabilisation (|ΔΔT_m_|= 5 K), hinting at a predominant non-specific interaction mechanism between [BMIM][BF_4_] and ubiquitin. This raises important questions about the predictive value of CSP in elucidating the nature of IL-protein interactions, suggesting that broader structural and environmental factors may play a decisive role (corroborating the previous pH and ionic strength results).

### Refining Interpretations of NMR Data Through SASA Ranked CSP Analysis

Thus, from the mutagenesis measurements, we did not see E16 to be extremely important for the IL mediated destabilisation despite having the highest CSP. Thus, we re-checked our NMR results keeping the accessibility information of different residues in mind. In NMR measurements, the CSP values indicate residue-specific perturbations in a protein caused by reaction conditions. Since, in our case, we used bulky ILs, the concern was whether the accessibility of IL also contributed to the NMR results. It is very likely that the impact of CSP values is scaled by the accessibility of the different residues to the IL. Thus, a high/low ΔCSP value might be a result of higher/lower accessibility. Moreover, small perturbations in the core can lead to significant destabilisation. To circumvent these problems, we performed a different type of analysis, SASA Ranked CSP Analysis (SRCA). We rearranged the ΔCSP values of the different residues according to their decreasing SASA (Fig. 6). We found that 7 out of 8 residues with ΔCSP > Mean + SD are among the 40% most surface-exposed residues, which is expected as they have a higher probability of interacting with the IL molecules. However, It’s also important to consider the buried residues, especially when discussing thermodynamic properties, as ILs are known to stabilize the unfolded state^22^ where these buried residues might gain exposure. Subsequently we arranged the amino acids in four different groups of hydrophobic, glycine, polar noncharged, and polar charged. Comparison the ΔCSP among these different groups on a rolling fashion across the SASA arranged residues yielded the hydrophobic residues and glycines to be majorly affected in presence of the IL. These observations established that hydrophobicity is the one of the key parameters. While the SRCA method provides valuable insights, it does have some limitations. Specifically, it does not account for potential neighbouring effects, where nearby residues can influence the extent of interaction with IL molecules. Therefore, SRCA should be used alongside other complementary methods to provide a more comprehensive understanding of IL-protein interactions.

**Figure 6.**
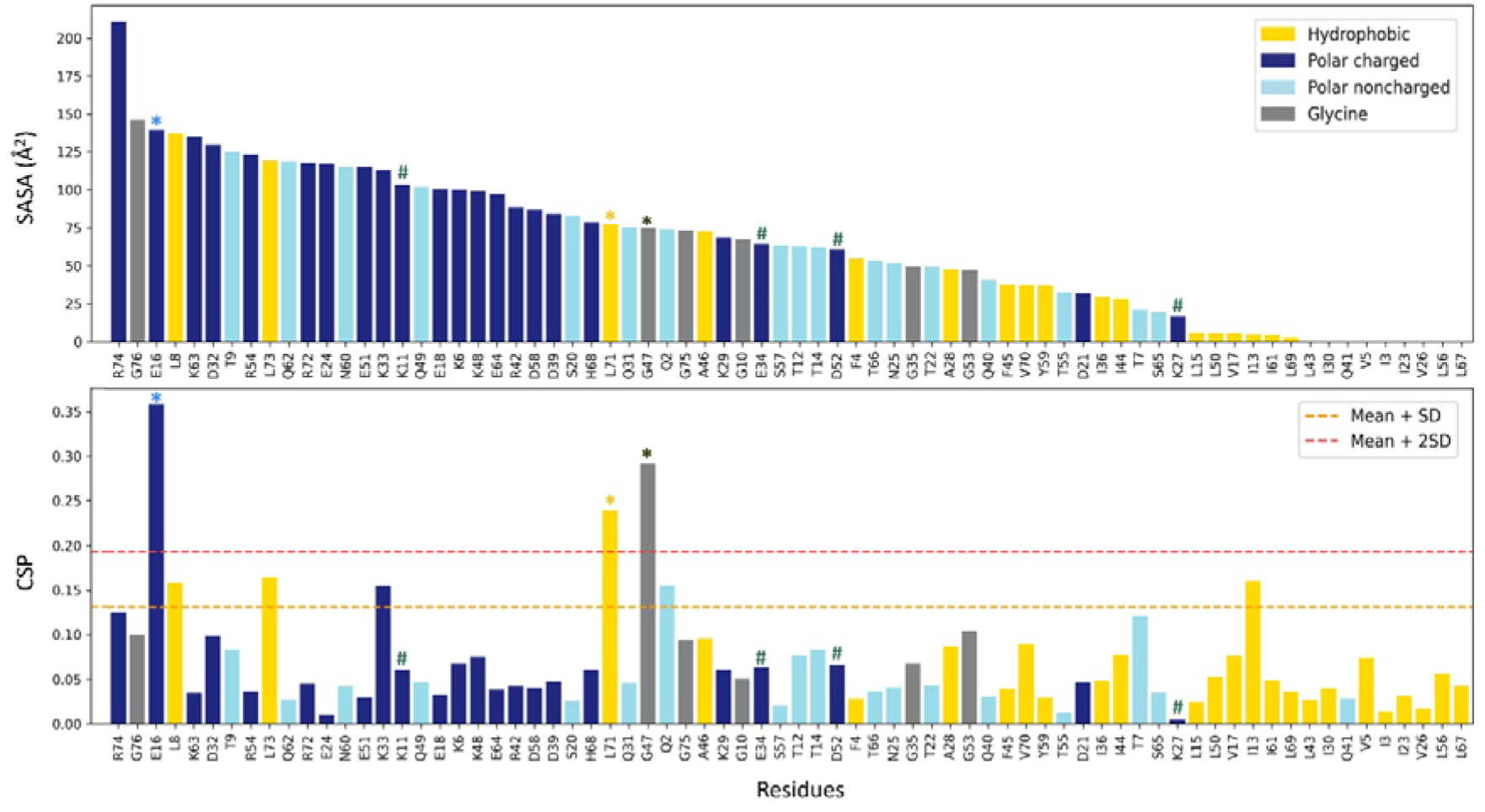
SASA Ranked CSP Analysis (SRCA) in Ubiquitin-[BMIM][BF_4_] interaction. The upper panel presents the SASA values for ubiquitin residues, sorted in descending order, while the lower panel displays the CSP values, indicating shifts due to interaction with [BMIM][BF4], to compare interaction propensities across residue types—hydrophobic, glycine, polar noncharged, and polar charged. Residues marked with an asterisk (*) and hashtag (#) are single mutants and quartet mutant respectively.

**Figure 7.**
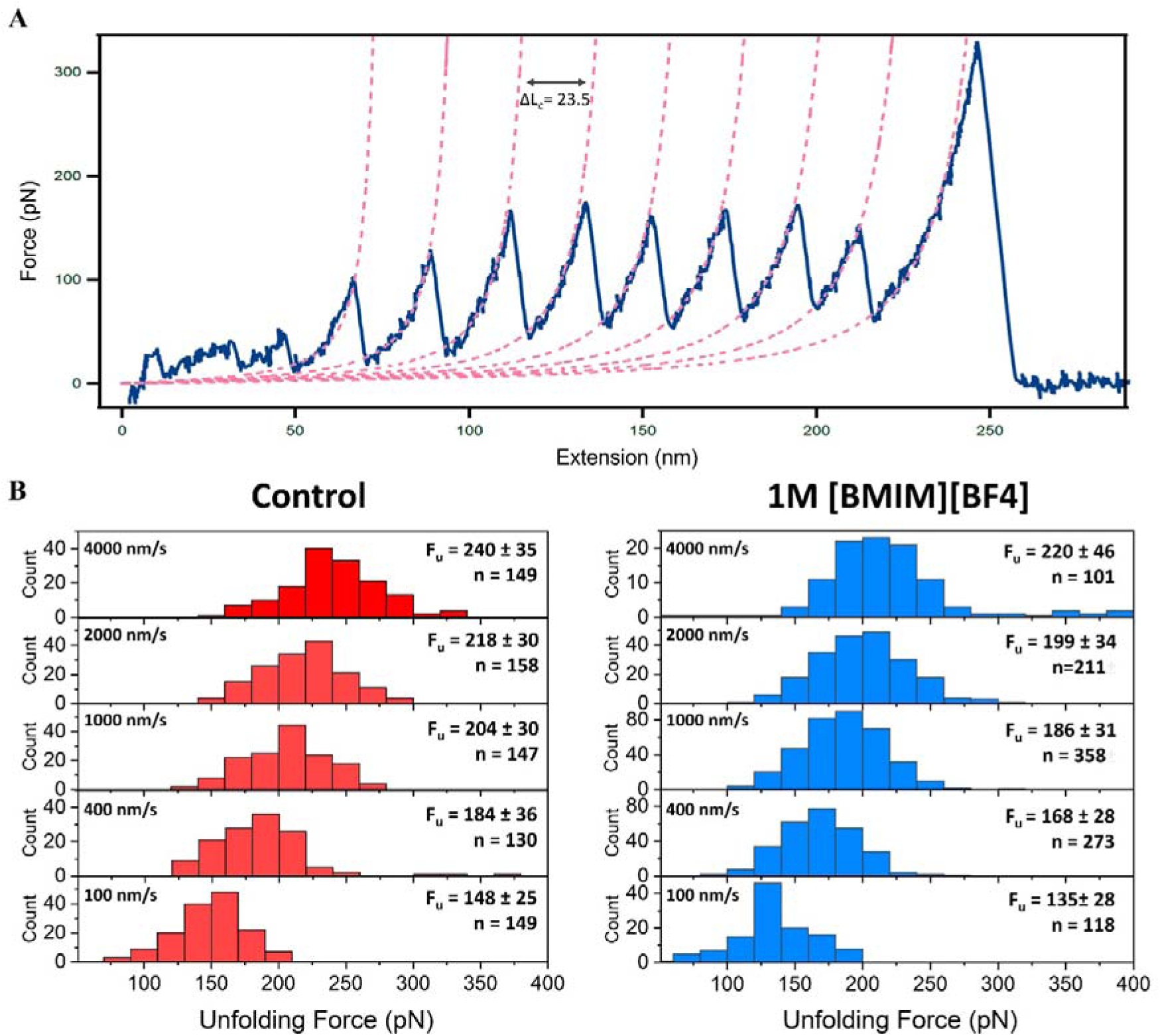
Mechanical stability of ubiquitin decreases in the presence of 1M [BMIM][BF_4_]. A) Representative FX unfolding trace of SpyTag003-(ubiquitin)_9_ showing a characteristic sawtooth pattern (blue solid line) with an unfolding contour length ΔL_c_ of ∼23.5 nm. The FX traces are fit with the WLC model of polymer elasticity (pink dashed lines), pulling speed – 400 nm/s. B) Histograms representing the unfolding forces at different pulling speeds (100 nm/s, 400 nm/s, 1000 nm/s, 2000 nm/s, 4000 nm/s) on the left in the absence (control) and on the right in the presence of [BMIM][BF4].

### [BMIM][BF_4_] makes the Protein Folding Energy Landscape shallower

Finally, we tried to correlate our findings by examining ubiquitin’s mechanical stability under IL influence via SMFS. Mechanical stability represents kinetic stability along the mechanical unfolding pathway, an unexplored aspect of protein behaviour in IL environments. Our SMFS findings reveal that [BMIM][BF4] notably decreases the force required for ubiquitin unfolding, suggesting ILs lower the kinetic barriers. This complements our thermodynamic observations, indicating a coherent effect of ILs on both the thermodynamic and mechanical stability of ubiquitin. Notably, Monte Carlo simulations revealed that the structural transition state, and thereby the protein’s flexibility, remains unchanged. SMFS results reveal that [BMIM][BF4] decreases the depth of the energy landscape of ubiquitin, lowering energy barriers for unfolding and suggesting a streamlined kinetic pathway while maintaining structural transition states as confirmed by Monte Carlo simulations. We observed a negligible change in the contour length (Fig. S5). This result aligns with experimental and computational observations. For instance, molecular dynamics (MD) simulations have demonstrated only a slight decrease in RMSD and a slight increase in the radius of gyration for α-lactalbumin^37^, or slight changes during SAXS-based studies for HSA and Cytochrome C^30^ in [BMIM][Cl]. However, there exist contrasting results as well, where significant changes have been observed, such as 6M [BMIM][NO3] causing notable conformational changes in lysozyme^32^. We proposed a simple model for the folding energy landscape to illustrate the impact of ILs on ubiquitin’s stability (Fig. 9). The model depicts a shallower energy profile in the presence of ILs, facilitating a faster transition from the destabilised native (N) state to the stabilised unfolded (U) state. This is supported by our observation of approximately a 10% reduction in near-UV CD spectra (Fig. S6) of the native state with ILs, indicating destabilisation. The presence of ILs reduces aggregation propensity, and a previous FCS ^30^ study has demonstrated that introducing an imidazolium-based IL ([PMIM][Br]) can stabilise the unfolded state, suggesting a stabilisation of the unfolded state.

**Figure 8.**
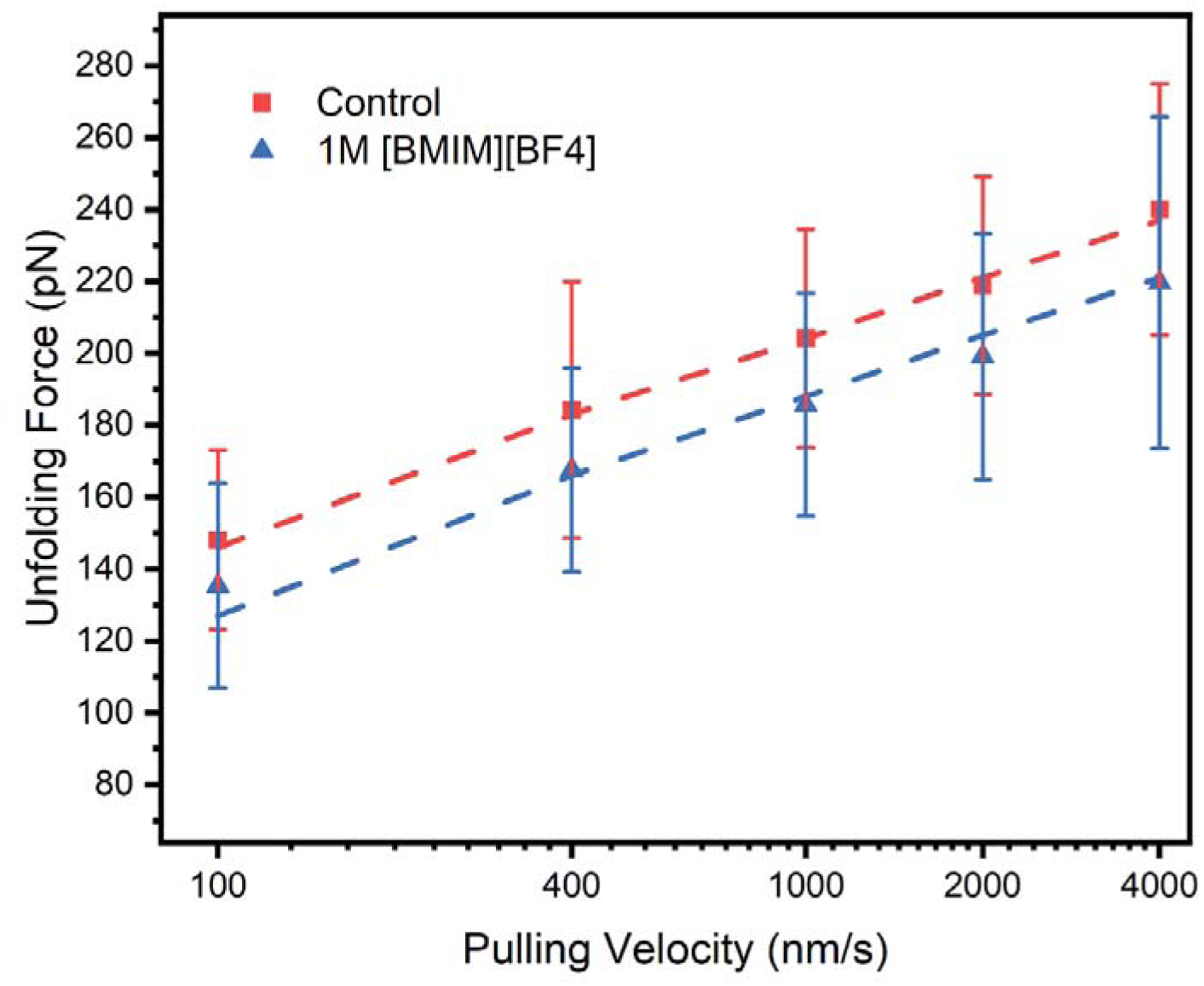
The spontaneous rate of unfolding of ubiquitin increases in the presence of 1M [BMIM][BF_4_]. The plot of the unfolding force vs pulling speed for ubiquitin in the absence (control) and presence of [BMIM][BF_4_], fit (dashed lines) was obtained from Monte Carlo simulations. The distance to the transition state remained the same.

**Figure 9.**
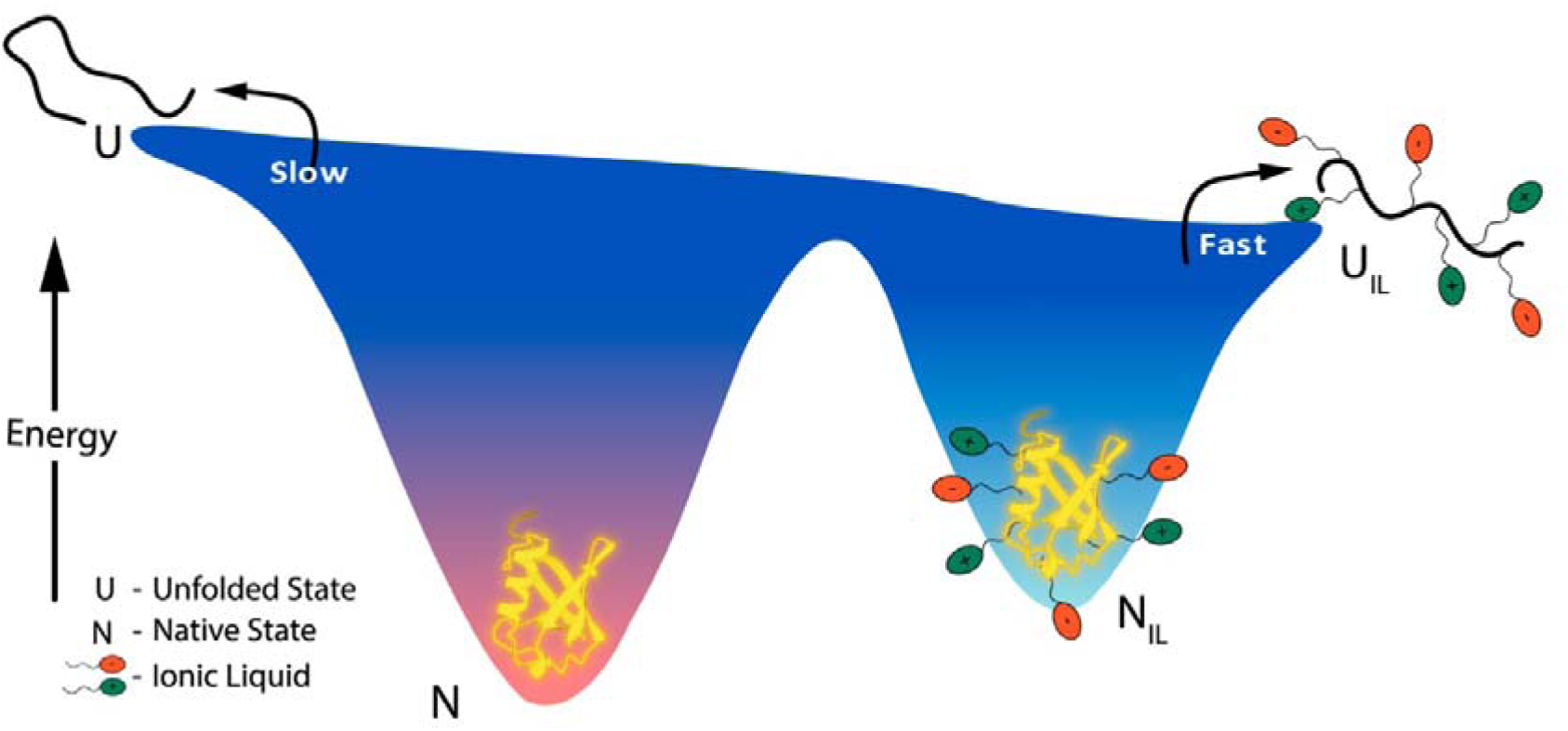
Impact of [BMIM][BF4] on the energy landscape of ubiquitin. The landscape depicts a shallower energy profile in the presence of ILs, facilitating a faster transition from the native (N) to the unfolded state (U). The ILs interact with both the native and unfolded forms, stabilising the unfolded state to prevent aggregation and destabilising the native state, as observed in the reduction in near-UV CD spectra.

## Conclusion

In this study, we have elucidated the impact of aqueous imidazolium-based ionic liquids (ILs) on the thermodynamic and kinetic stability of ubiquitin, employing techniques such as circular dichroism (CD) spectroscopy, nuclear magnetic resonance (NMR), and single-molecule force spectroscopy (SMFS). Our findings corroborate that hydrophobic interactions are the critical factor influencing the stability of ubiquitin. Importantly, we demonstrated that variations in IL cations and anions significantly influence protein-IL interactions. Cations with longer alkyl chains and greater hydrophobicity ([BMIM]^+^ > [BMPyr]^+^ > [EMIM]^+^) led to increased destabilization of ubiquitin, highlighting the role of hydrophobic interactions in modulating protein stability. Similarly, anions contributed to destabilization following the Hofmeister series order [BF_]_ > [MeSO_]_ > [Cl]_ with some deviations, indicating that the choice of anion affects the extent of interactions with the protein. The extent of destabilisation could be modulated through adjustments in solvent conditions like ionic strength and pH. Further, our detailed CSP based NMR analysis, identified specific residues of ubiquitin affected by [BMIM][BF4]. However, targeted mutagenesis of these residues did not markedly alter protein destabilisation. On incorporating SASA into our empirical evaluation, we found a predominantly global effect of [BMIM][BF4] on ubiquitin rather than localized interactions. This observation is crucial as it suggests that ILs do not single-handedly dictate stability changes through specific residues but rather through broader hydrophobic interactions.

Moreover, SMFS experiments revealed that ILs lower the unfolding barrier of ubiquitin without modifying the transition state structures, providing deeper insights into the protein folding dynamics under the influence of ILs. These results highlight how ILs energetically and kinetically favour the unfolded state, which could significantly influence protein-protein or protein-substrate interactions. To further advance our understanding and harness the full potential of ILs in biotechnology and pharmaceutical applications, it is imperative to expand the scope of SMFS studies in different ILs. Ultimately, the fundamental insights gained from this study can be instrumental in devising translational applications that exploit IL-mediated modulations of protein behaviour for enhanced therapeutic and industrial outcomes.

## Abbreviations

AFM: Atomic Force Microscope
APDMS: 3-Aminopropyl(diethoxy)methylsilane
BMIM: 1-Butyl-3-methylimidazolium tetrafluoroborate
BMPyr: 1-Butyl-1-methylpyrrolidinium
CD: circular dichroism
EMIM: 1-Ethyl-3-methylimidazolium tetrafluoroborate
IL: ionic liquid
IPTG: isopropyl β-D-1-thiogalactopyranoside
LB: Lysogeny Broth

## Acknowledgment

We acknowledge the generous help from, Dr. Arpan Dey, Ms. Mamata Kallianpur, Mr. Rishi Verma, and Mr. Satyanarayan at the different stages of this project. We thank the TIFR and DAE (Department of Atomic Energy, India) for financial support under Project No. 12-R&DTFR-5.10-0100.

