## Supplementary Information for "Ionic Liquid-Induced Modulation of Ubiquitin Stability: The Dominant Role of Hydrophobic Interactions"

**Protein Sequences:**

1. UF45W :

MRGSHHHHHHGSMQIFVKTLTGKTITLEVEPSDTIENVKAKIQDKEGIPPDQQRLIWAGKQLEDGRTLSDYNIQKESTLHLVLRLRGGRS-

2. Spytag003-(Ubq)9:

MGRGVPHIVMVDAYKRYKSGGGSMQIFVKTLTGKTITLEVEPSDTIENVKAKIQDKEGIPPDQQRLIFAGKQLEDGRTLSDYNIQKESTLHLVLRLRGGMQIFVKTLTGKTITLEVEPSDTIENVKAKIQDKEGIPPDQQRLIFAGKQLEDGRTLSDYNIQKESTLHLVLRLRGGMQIFVKTLTGKTITLEVEPSDTIENVKAKIQDKEGIPPDQQRLIFAGKQLEDGRTLSDYNIQKESTLHLVLRLRGGMQIFVKTLTGKTITLEVEPSDTIENVKAKIQDKEGIPPDQQRLIFAGKQLEDGRTLSDYNIQKESTLHLVLRLRGGMQIFVKTLTGKTITLEVEPSDTIENVKAKIQDKEGIPPDQQRLIFAGKQLEDGRTLSDYNIQKESTLHLVLRLRGGMQIFVKTLTGKTITLEVEPSDTIENVKAKIQDKEGIPPDQQRLIFAGKQLEDGRTLSDYNIQKESTLHLVLRLRGGMQIFVKTLTGKTITLEVEPSDTIENVKAKIQDKEGIPPDQQRLIFAGKQLEDGRTLSDYNIQKESTLHLVLRLRGGMQIFVKTLTGKTITLEVEPSDTIENVKAKIQDKEGIPPDQQRLIFAGKQLEDGRTLSDYNIQKESTLHLVLRLRGGMQIFVKTLTGKTITLEVEPSDTIENVKAKIQDKEGIPPDQQRLIFAGKQLEDGRTLSDYNIQKESTLHLVLRLRGGCCGGTGSGSHHHHHH-

3. ELP_120_-SpyCatcher003:

MCVPGEGVPGVGVPGVGVPGVGVPGVGVPGAGVPGAGVPGGGVPGGGVPGEGVPGEGVPGVGVPGVGVPGVGVPGVGVPGAGVPGAGVPGGGVPGGGVPGEGVPGEGVPGVGVPGVGVPGVGVPGVGVPGAGVPGAGVPGGGVPGGGVPGEGVPGEGVPGVGVPGVGVPGVGVPGVGVPGAGVPGAGVPGGGVPGGGVPGEGVPGEGVPGVGVPGVGVPGVGVPGVGVPGAGVPGAGVPGGGVPGGGVPGEGVPGEGVPGVGVPGVGVPGVGVPGVGVPGAGVPGAGVPGGGVPGGGVPGEGLPETGGSAMVTTLSGLSGEQGPSGDMTTEEDSATHIKFSKRDEDGRELAGATMELRDSSGKTISTWISDGHVKDFYLYPGKYTFVETAAPDGYEVATPIEFTVNEDGQVTVDGEATEGDAHTGHHHHHH-

**Primers used:**

| **Primer Name** | **Primer Sequence** |
| --- | --- |
| UF_E16A_Fwd | CCATAACTCTGGCGGTTGAACCATCCGATACCATCGAAAACGT |
| UF_E16A_Rvs | CAACCGCCAGAGTTATGGTTTTACCGGTTAACGTCTTGACG |
| Ubq_G47A_Fwd | GCCGCGAAGCAGCTCGAGGACGGTAGAACGCTGTCTGATT |
| Ubq_G47A_Rvs | CGAGCTGCTTCGCGGCCCAGATCAATCTTTGTTGATCAGG |
| Ubq_L71A_Fwd | CTGGTCGCGAGACTAAGAGGTGGTAGATCTTGAGGTACCCCGG |
| Ubq_L71A_Rvs | CTTAGTCTCGCGACCAGATGTAAGGTCGACTCCTTCTGAATG |
| Ubq9_Fwd | GCGTATCACGAGGCCCTTTCGTCT |
| Ubq9_Rvs | CGACCCGGGGTACCTAAGCAACAAC |
| ELP_Bam_ins_fwd | GGATCCGGCTAACTCGAGTAAGATCCGGCTGCTAACAAAGC |
| ELP_Bam_ins_rvs | CTCGAGTTAGCCGGATCCGCCGGTTTCCGGCAGACCTTCAC |
| Spycat_BamHI_fwd | CCTGTATTTTCAGGGATCCGCCATGGTAACCACCTTATCAG |
| Spycat_his_xho_rvs | CCCTCGAGTTAATGGTGATGATGGTGATGTCCAGTATGAGCGTCACCTTCAGTTGCTTC |
| Spycat_C49S_fwd | GAGTTGCGTGATTCCTCTGGTAAAACTATTAGTACATGGATTTCAG |
| Spycat_C49S_rvs | GAGGAATCACGCAACTCCATAGTTGCACCAGCTAACTCAC |

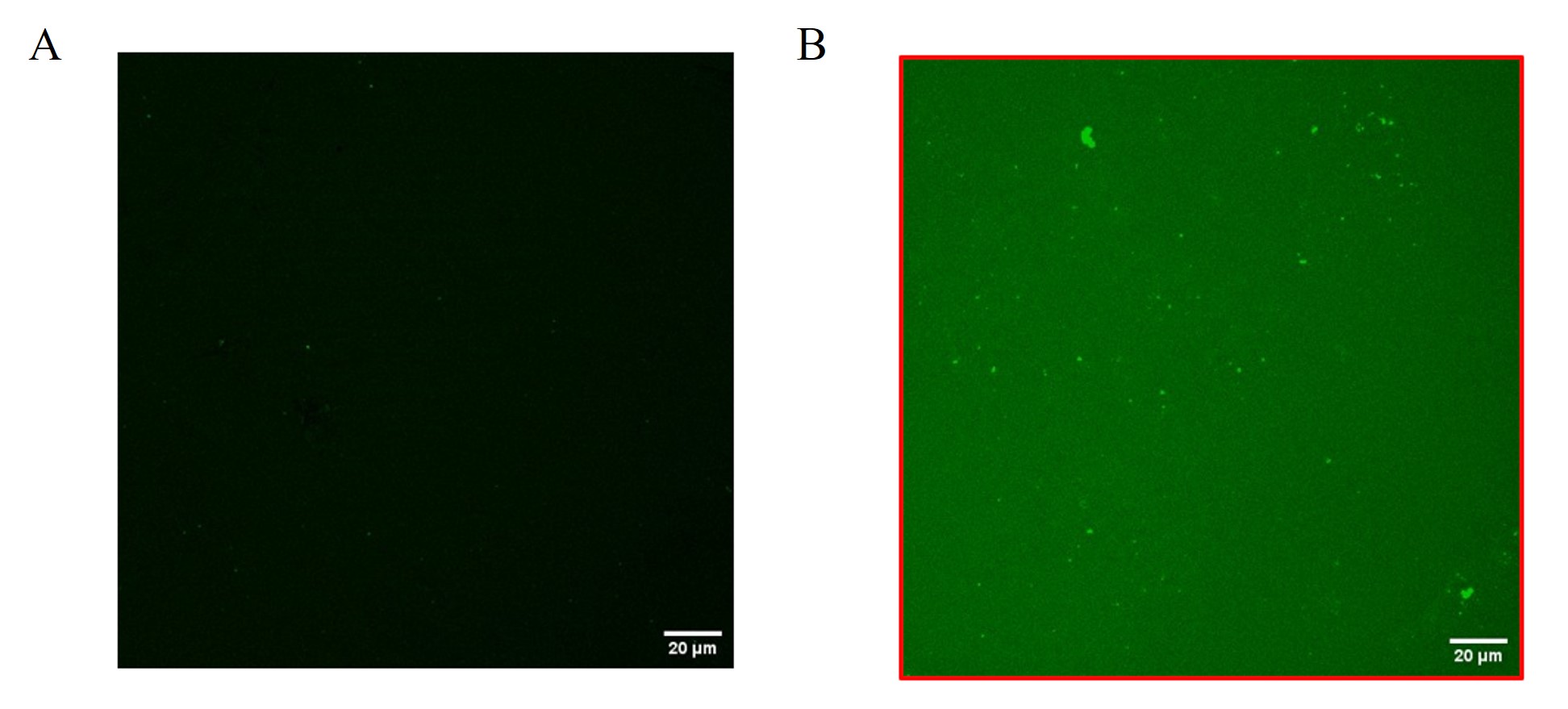

**Fig S1.** Confocal microscopy images of coverslips after incubating with SpyTag003-mKate2 and washing. a) Control coverslip without immobilisation of SpyCatcher003, b) Coverslip with SpyCatcher003 immobilised. Excitation – 543 nm, emission was collected 556-700 nm.

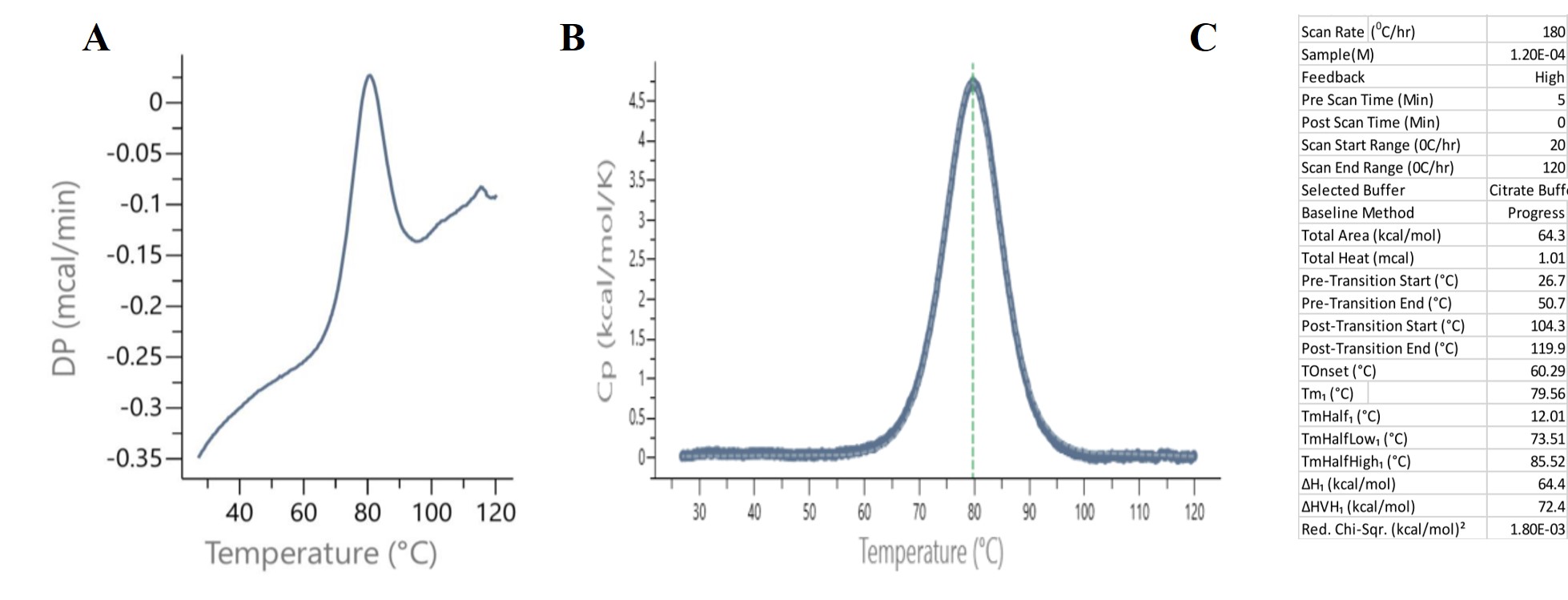
 **Fig S2.** Differential Scanning Calorimetry (DSC) analysis of ubiquitin variant UF45W at pH 3.5. A) Raw DSC Thermogram (Δp vs. Temperature), highlighting the thermal denaturation process of ubiquitin, with the peak corresponding to the melting temperature (T_m_) of the protein. B) Fitted DSC Thermogram (Cp vs. Temperature). C) Parameters used during the experiments and quantities obtained after the analysis.

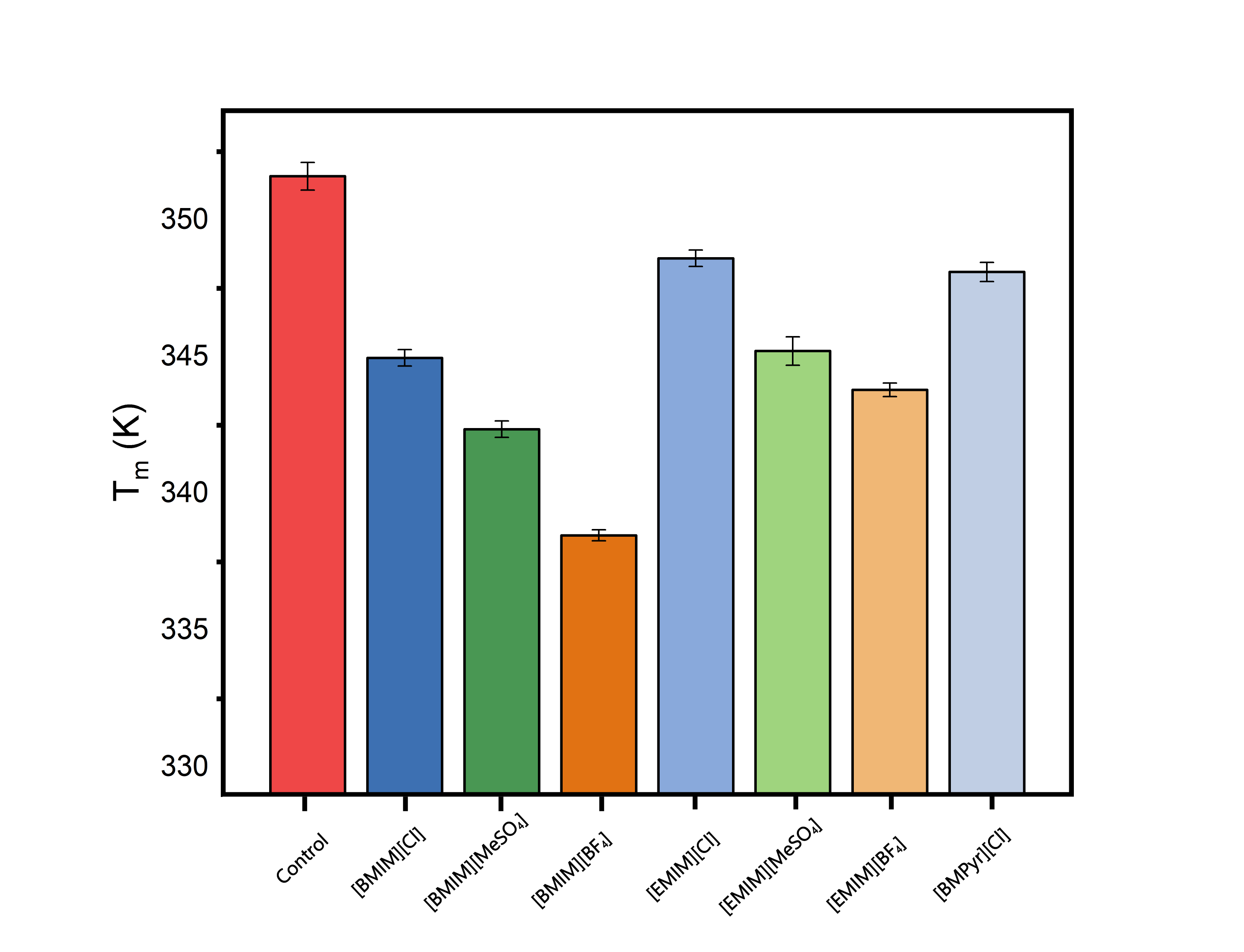

**Fig S3**. Analysis of ubiquitin variant UF45W thermal stability in ILs. Bar graph representing Tm analysis in different ionic liquids at 0.5 M concentration, demonstrating the influence of ionic environments on protein stability.

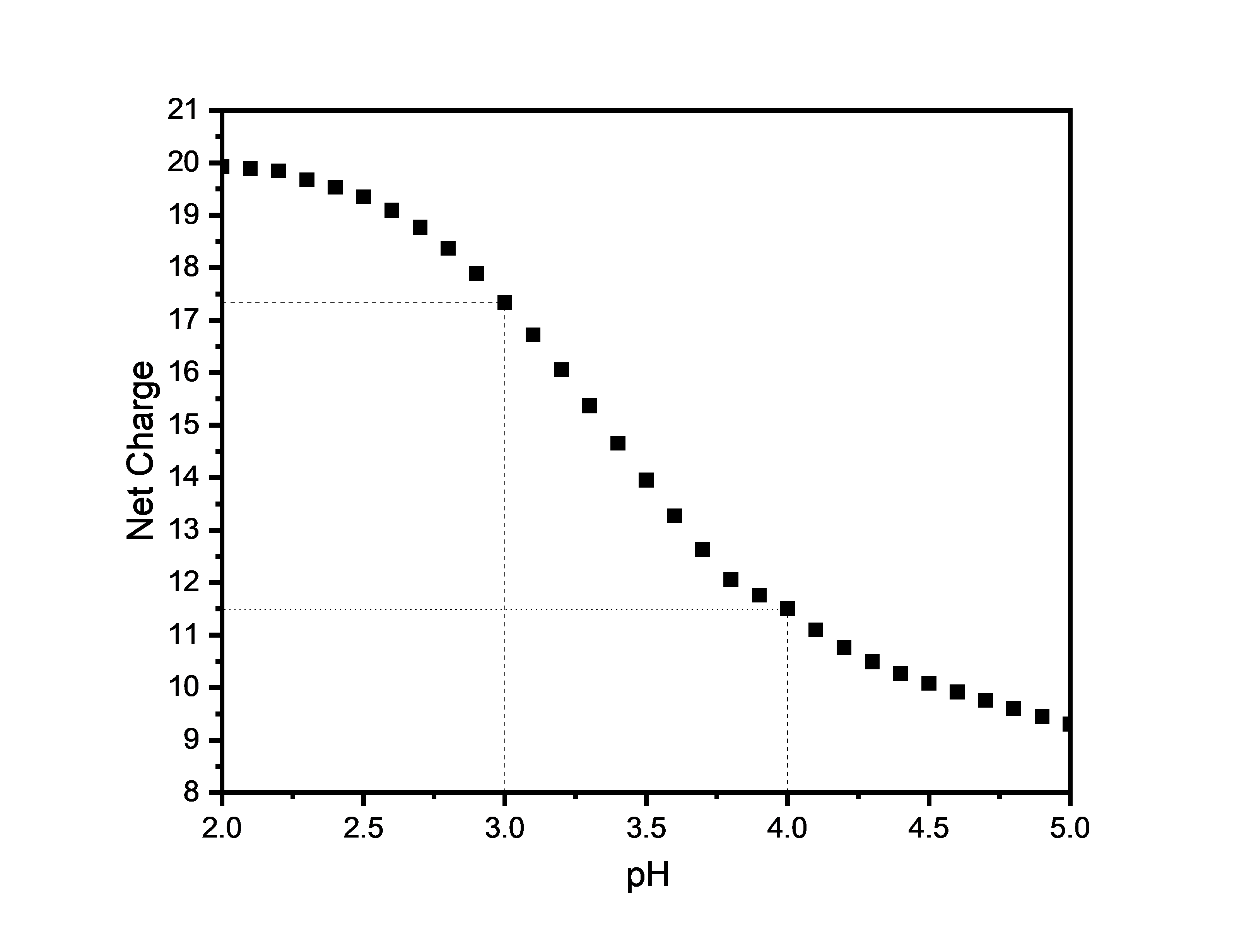

**Fig S4.** The net charge of ubiquitin variant UF45W under various pH conditions, as calculated by the DelPhi Webserver utilizing the DelPhi-PKA tool.

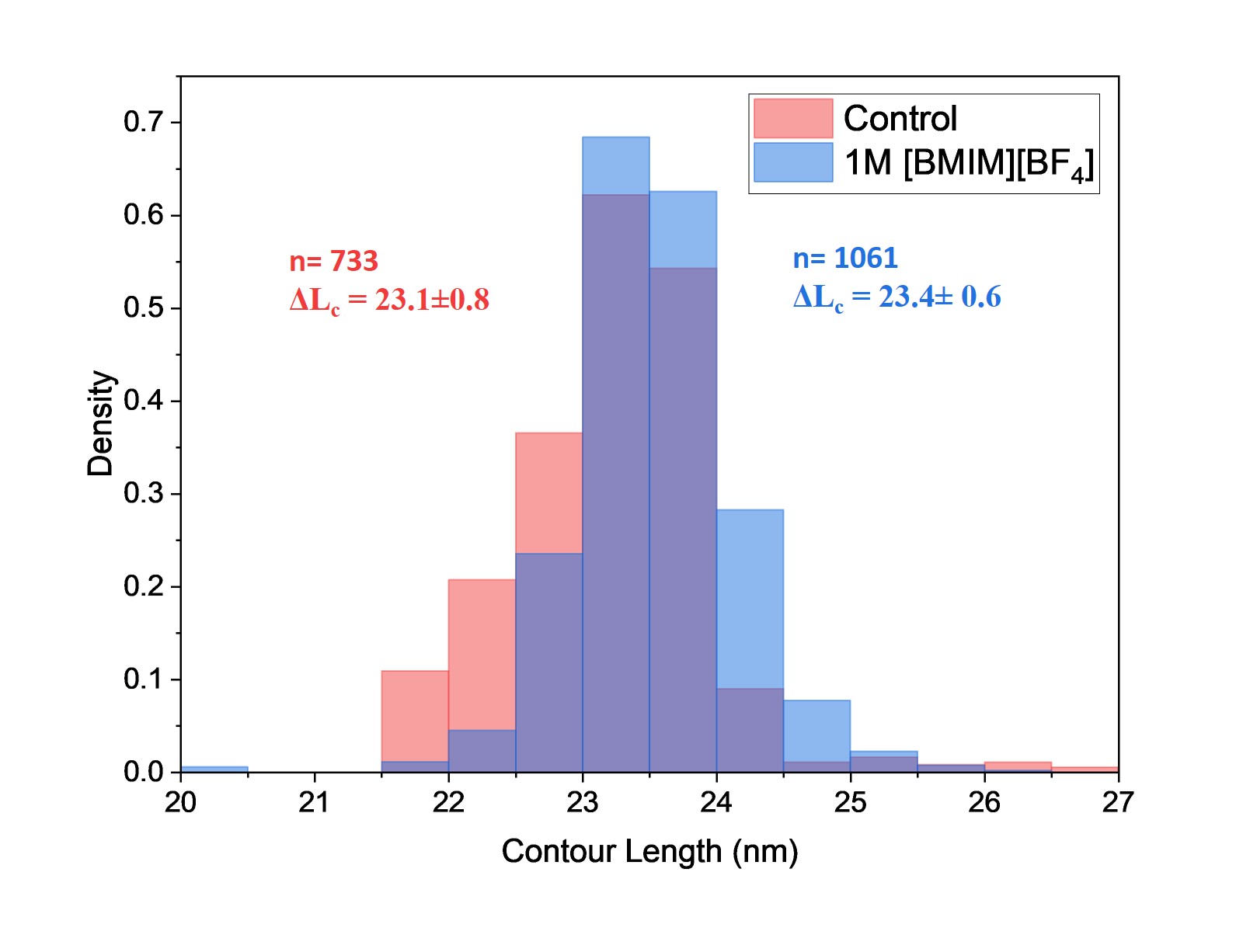

**Fig. S5.** Contour length of ubiquitin remains the same in presence of 1M [BMIM][BF_4_].

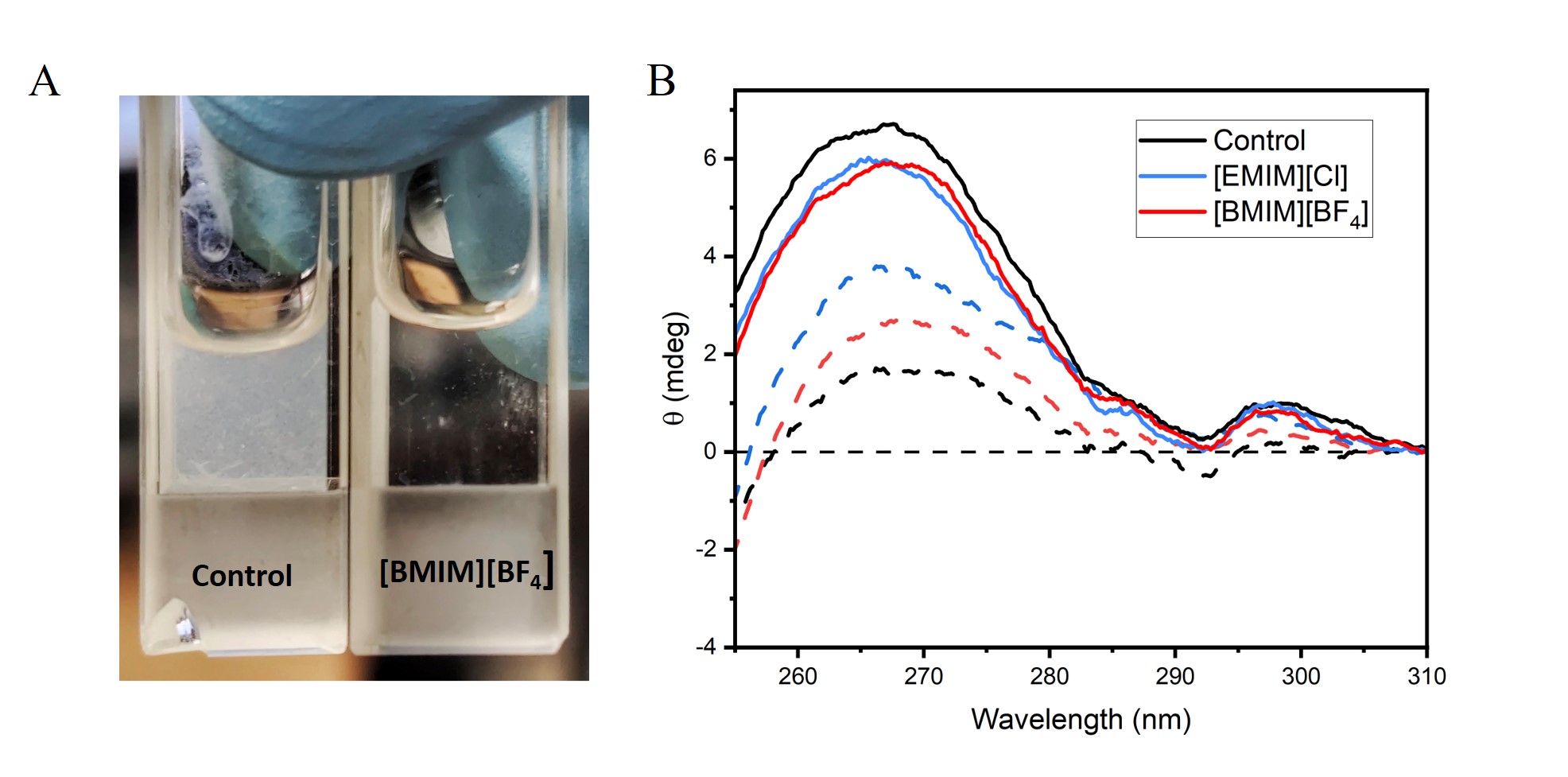

**Fig. S6.** ILs reduces ubiquitin aggregation propensity. A) Image of cuvettes comparing UF45W solutions after a thermal denaturation cycle. The left cuvette contains the control sample, while the right cuvette shows the sample in 1M [BMIM][BF4], pH 4, highlighting the reduced aggregation - both in quantity and size - in the presence of IL. B) Near-UV CD spectra of UF45W before thermal denaturation (solid lines), and post denaturation (dashed lines). The presence of IL is shown to enhance the refolding fraction, evidenced by increased ellipticity in the CD spectra after denaturation as compared to the control (The concentration of ILs was 0.5M, pH 3).

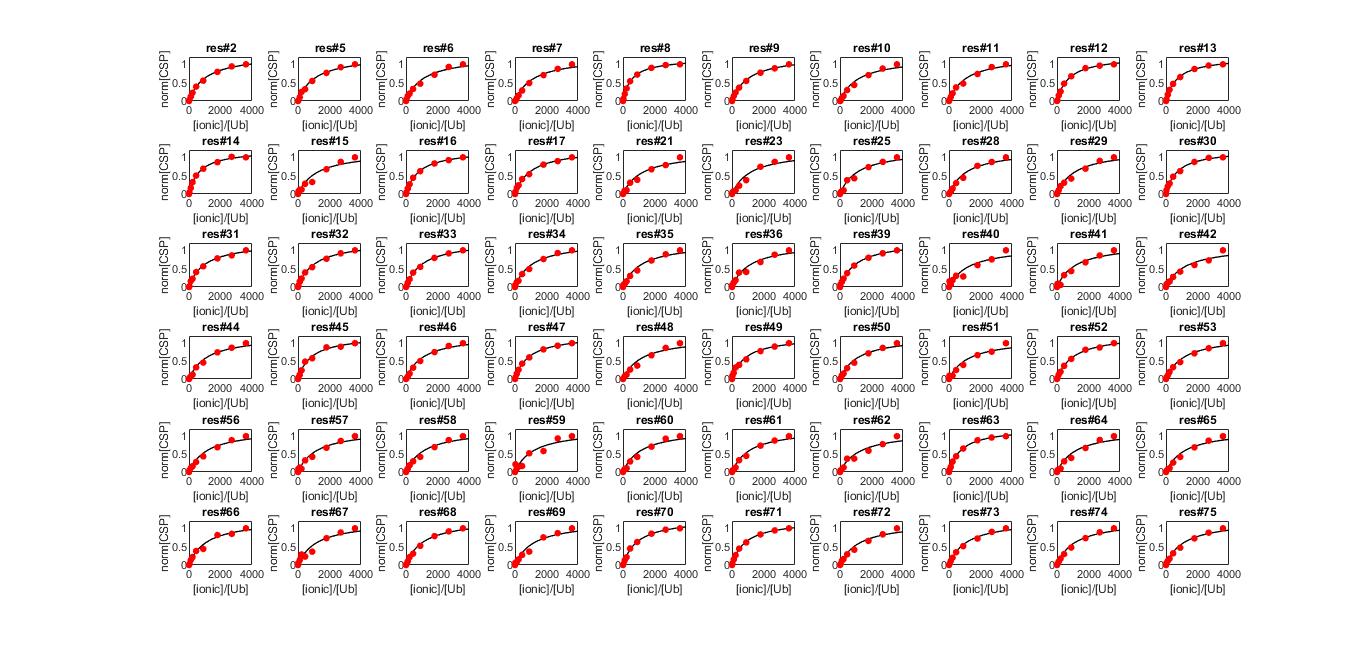

**Fig S7.** The fit of NMR titration data of Ubiquitin with BMIM-BF4. CSPs calculated for each residue at every titration point were normalized with the highest value and fitted into a 1:1 protein-ligand model using the equation described (Materials and methods) to evaluate per residue K_d_. Average K_d_ = 311 ± 8 mM.

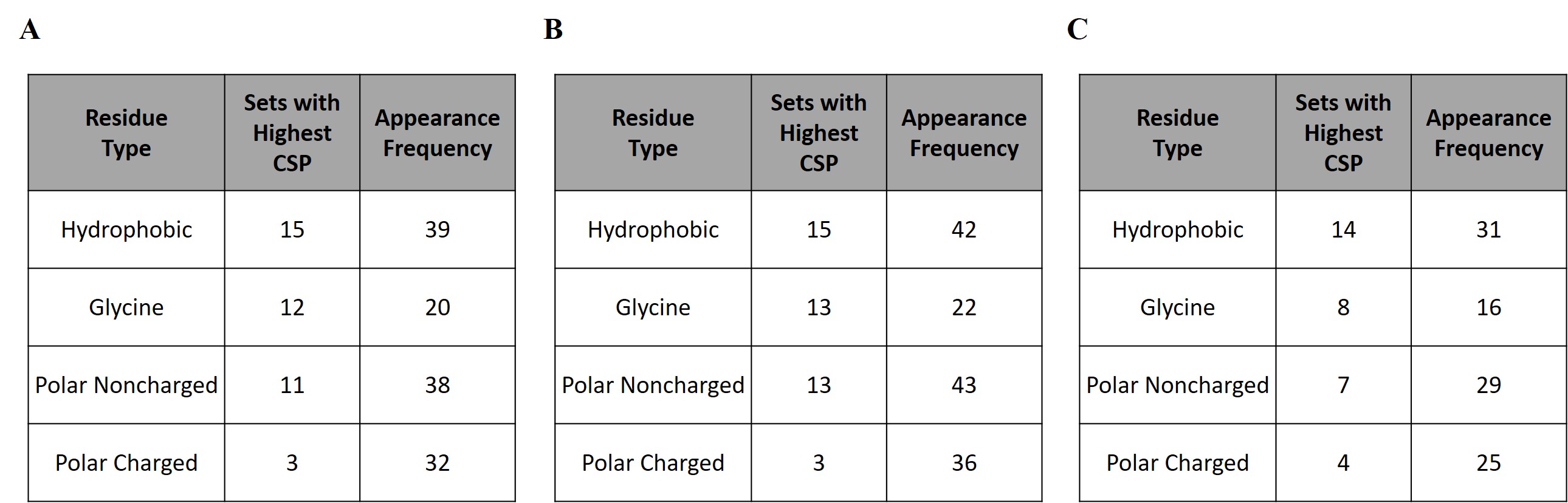
**Table S1.** Tables consisting the SASA Ranked CSP Analysis (SRCA) by examining moving sets of A) 5 (n=41) B) 6 (n=44) and C) 4 (n=31) consecutive residues. After analysing each set, the frequency of each category (hydrophobic, glycine, polar non-charged, and polar charged) with highest CSP and the appearance frequency were calculated.

**Script-1**

import freesasa

from Bio.PDB import PDBParser

import csv

\# Define residue categories

hydrophobic = ['A', 'V', 'I', 'L', 'M', 'F', 'Y', 'W']

polar_uncharged = ['S', 'T', 'N', 'Q']

polar_charged = ['R', 'H', 'K', 'D', 'E']

glycine = ['G'] # Glycine

\# Define a dictionary for converting three-letter to one-letter residue codes

three_to_one_map = {

'ALA': 'A', 'CYS': 'C', 'ASP': 'D', 'GLU': 'E', 'PHE': 'F',

'GLY': 'G', 'HIS': 'H', 'ILE': 'I', 'LYS': 'K', 'LEU': 'L',

'MET': 'M', 'ASN': 'N', 'PRO': 'P', 'GLN': 'Q', 'ARG': 'R',

'SER': 'S', 'THR': 'T', 'VAL': 'V', 'TRP': 'W', 'TYR': 'Y'

}

\# Function to categorize residues

def categorize_residue(residue_code):

if residue_code in hydrophobic:

return 'Hydrophobic'

elif residue_code in polar_uncharged:

return 'Polar noncharged'

elif residue_code in polar_charged:

return 'Polar charged'

elif residue_code in glycine:

return 'Glycine'

else:

return 'Unknown'

\# Load the structure from a PDB file

pdb_file = '1ubq.pdb'

structure = freesasa.Structure(pdb_file)

\# Calculate the SASA

result = freesasa.calc(structure)

\# Use Bio.PDB to parse residue names

parser = PDBParser(QUIET=True)

bio_structure = parser.get_structure('1UBQ', pdb_file)

\# Get SASA for all residues

residue_sasa_dict = result.residueAreas()

sasa_dict_chain = residue_sasa_dict['A'] # Only chain A

\# Prepare data for CSV

csv_data = [['Residue', 'SASA', 'Category']]

\# Collect residue data

for model in bio_structure:

for chain in model:

if chain.id == 'A':

for residue in chain:

residue_id = residue.id[1]

residue_name = residue.get_resname()

one_letter_code = three_to_one_map.get(residue_name, '?')

\# Retrieve SASA for the residue

sasa_key = str(residue_id) # Use residue number as string

if sasa_key in sasa_dict_chain:

sasa = sasa_dict_chain[sasa_key].total

category = categorize_residue(one_letter_code) # Determine category

\# Add Residue name with number (e.g., M1), SASA, and category to CSV data

csv_data.append([f"{one_letter_code}{residue_id}", f"{sasa:.2f}", category])

\# Write data to CSV file

output_file = 'residue_sasa.csv'

with open(output_file, 'w', newline='') as csvfile:

csv_writer = csv.writer(csvfile)

csv_writer.writerows(csv_data)

print(f"Residue-wise SASA data has been written to {output_file}.")

**Script-2**

import pandas as pd

\# Load the data

df = pd.read_csv('movingCSP.csv')

\# Initialize an empty list to store the results

results = []

\# Define your new parameters

window_size = 5 # change this to your preferred window size

min_residue_types = 3 # change this to your preferred minimum number of different types of residues

\# Create an empty dictionary to store the counts

residue_type_counts = {'Hydrophobic': 0, 'Polar charged': 0, 'Polar noncharged': 0, 'Glycine': 0}

\# Loop over each set of consecutive residues with size = window_size

for i in range(len(df) - (window_size - 1)):

subset = df.iloc[i:i+window_size]

\# Check if the subset contains at least min_residue_types different types of residues

if subset['Type of residue'].nunique() >= min_residue_types:

\# Find the residue with the highest CSP

max_csp_residue = subset.loc[subset['CSPs'].idxmax()]

\# Append the result to the list

results.append((max_csp_residue['Residue'], max_csp_residue['Type of residue'], max_csp_residue['CSPs']))

\# Count the unique residue types in the chunk and update the dictionary

unique_types = subset['Type of residue'].unique()

for residue_type in unique_types:

residue_type_counts[residue_type] += 1

\# Convert the results to a DataFrame

results_df = pd.DataFrame(results, columns=['Residue', 'Type of residue', 'CSP'])

\# Print the results

print(results_df)

print(residue_type_counts)

\# Count the number of times each type of residue appears in the results

counts = results_df['Type of residue'].value_counts()

\# Print the counts

print(counts)
